# Microtubule severing enzymes oligomerization and allostery: a tale of two domains

**DOI:** 10.1101/2022.07.26.501617

**Authors:** Amanda C. Macke, Maria S. Kelly, Rohith Anand Varikoti, Sarah Mullen, Daniel Groves, Clare Forbes, Ruxandra I. Dima

## Abstract

Severing proteins are nanomachines from the AAA+ (ATPases associated with various cellular activities) superfamily whose function is to remodel the largest cellular filaments, microtubules. The standard AAA+ machines adopt hexameric ring structures for functional reasons, while being primarily monomeric in the absence of the nucleotide. Both major severing proteins, katanin and spastin, are believed to follow this trend. However, studies proposed that they populate lower-order oligomers in the presence of co-factors, which are functionally relevant. Our simulations show that the preferred oligomeric assembly is dependent on the binding partners, and on the type of severing protein. Essential dynamics analysis predicts that the stability of an oligomer is dependent on the strength of the interface between the helical bundle domain (HBD) of a monomer and the convex face of the nucleotide binding domain (NBD) of a neighboring monomer. Hot spots analysis found that the region consisting of the HBD tip and the C-terminal (CT) helix is the only common element between the allosteric networks responding to nucleotide, substrate, and inter-monomer binding. Clustering analysis indicates the existence of multiple pathways for the transition between the secondary structure of the HBD tip in monomers and the structure(s) it adopts in oligomers.

## INTRODUCTION

A special class of AAA+ (ATPases Associated with diverse cellular Activities) molecular machines, which convert chemical energy from ATP hydrolysis into mechanical force to assist essential cellular mechanisms, is represented by microtubule severing proteins. The main types of severing enzymes are katanin and spastin. These proteins are involved in the remodeling of the longest and stiffest assembly in the cell, the microtubule filament, during fundamental processes such as cellular division, neuronal growth, and ciliogenesis. While their original action was confined to cutting microtubules at locations far from their ends, recent studies^1^ revealed that severing proteins can also help amplify microtubule arrays and, at least in the case of katanin, catalyze the depolymerization of microtubules both in *vitro* and in *vivo* by acting on the plus-end of the filament.^2^ The ATPase region in both severing proteins is composed of two domains: the nucleotide-binding domain (NBD), which is common to all AAA+ protein family members, and the four–helix bundle domain (HBD). The NBD consists of highly conserved elements among AAA+ proteins: the Walker–A (WA), Walker–B (WB), Arg–finger (RR) motifs, and central pore loops 1, 2, and 3 (PL1, PL2, and PL3). The textbook behavior of AAA+ enzymes involved in protein remodeling is that, in the presence of ATP, monomers assemble into hexameric rings that form a narrow pore. This central pore, formed by NBD regions from each monomer, is functionally crucial as, during ATP-driven allosteric cycles, flexible channel loops from each NBD bind to the substrate and mediate the application of forces to promote remodeling of their substrates through unfolding or disassembly. In recent years, cryo-EM structures of severing proteins, solved in the presence of a minimal substrate (poly-Glu) peptides and nucleotides, revealed that these enzymes, similar to many other members of the AAA+ family such as ClpB/Hsp104, ClpA, ClpX, and ClpY, acquire non–planar arrangements of monomers in their hexameric structures resulting in “spiral” or “ring” conformations.^3–9^ The central pore loops PL1 and PL2 from each monomer form a double-helical staircase gripping the odd and even Glu residues of the substrate peptide, respectively. This arrangement is distinct from the one found in other AAA+ proteins where the central pore loops grip only every other residue of a substrate peptide,^10^ and is likely a signature of the tight grip that severing proteins exert on their substrate. Experimental studies also proposed that PL1 and PL2 in severing proteins are allosterically coupled to the ATP binding site and the oligomerization interfaces, which can provide a structural explanation for the tubulin substrate binding-induced katanin and spastin activation.^11,12^ The HBD region in severing proteins, while less well characterized from a functional point of view than the NBD, adopts a semi crescent structure covering the NBD of an adjacent monomer in the oligomeric species.^13^ Namely, in the hexameric states, the interior monomers (B to E) make canonical convex–to–concave AAA+ interactions with their nearest neighbors: the convex face of NBD from monomer i interacts with both the NBD and the HBD from monomer i-1, while the concave face of NBD from monomer i interacts only with the NBD of monomer i+1.^12,14^

In AAA+ proteins such as ClpA and ClpB, the non-planar structure of the hexameric assembly suggested a “hand-over-hand” substrate protein translocation mechanism driven by non concerted ATP hydrolysis in each monomer, comprising of the substrate protein (SP) grip and release by successive central channel loops of each AAA+ domain.^15^ The existence of a similar mechanism for the action of severing proteins is not yet established. ^12,14,16,17^ Several proposed mechanisms are: unfoldase, which is similar to the hand-over-hand action, wedge-like, or a combined unfoldase and wedge named death-spiral.^16,18–20^ Our previous studies showed that, while the unfoldase action is important,^21,22^ the wedge mechanism is most likely making a large contribution to the disassembly of a microtubule by severing proteins.^23–25^

An important functional aspect of severing proteins found in experimental studies is the instability in the hexamers upon ATP hydrolysis and/or disengagement from the substrate protein. This led to the proposal that the dynamic dissociation of the hexamer acts as a mechanism for disengaging severing proteins from microtubules to facilitate rapid and efficient processing of the filament^26^ and that severing proteins are non processive enzymes. ^13^ This is supported by recent experiments for another AAA+ protein from the same meiotic clade as severing proteins, ClpB, which found that this machine might specifically target unfolded portions of its typical substrates (protein aggregates) for translocation followed by the release of the substrate upon encountering a folded domain. This alternation between processive translocation on unfolded regions and then release would in turn make the overall translocation by ClpB non-processive.^27^ Interestingly, the disassembly of the hexameric state into lower order oligomers also occurred in our recent simulations of the action of spastin on microtubule filaments, especially when the interactions between the severing machine and the microtubule surface reach levels comparable to the interactions between kinesin-1 motors and microtubules.^22^

Another fascinating aspect of microtubule severing enzymes is that, unlike other machines from the AAA+ family, which require only ATP binding to reach their oligomeric functional configuration, they are believed to form higher-order oligomers (hexamers) only if their substrate, a MT filament or, at the minimum, the C-terminal tail (CTT) from a tubulin subunit, is present as well.^28^ For example, for ClpB sedimentation studies found that, in the absence of nucleotides, ClpB from E.*coli* undergoes reversible self-association that involves protein concentration-dependent populations of monomers, higher-order oligomers (heptamers or hexamers) and intermediate-size oligomers, most likely dimers. The presence of nucleotides leads to the stabilization of ClpB hexamers, at the expense of any lower-order oligomers.^29^ Studies investigating the oligomerization process in severing proteins pointed towards multi-step mechanisms, which are likely distinct for these two machines. For example, biochemistry studies proposed that spastin binds to a microtubule in monomeric form via electrostatic interactions, followed by diffusion on the microtubule surface leading to the formation of lower-order oligomers, likely dimers.^30^ These serve as nucleated assembly for the rapid formation of hexameric states. The human forms of spastin and katanin both bind to microtubules and move along the filament diffusively with similar velocities, without the need for ATP hydrolysis.^30^ Other studies^13^ showed that C.*elegans* katanin in the absence of ATP and the substrate oligomerizes into a mixture of dimers and trimers. At the same time, for H.*sapiens* katanin the stable forms are dimers and monomers. ^13^ The A.*thaliana* katanin was shown to oligomerize into trimers, with the MIT domains playing a role in the process.^31^ Finally, studies^14,32^ suggested that katanin p60 shifts from monomeric to hexameric fractions in a concentration-dependent manner and, at high concentrations, both ATP and the microtubule filament are dispensable for the formation of the hexamer. These results suggest that the assembly of the ATPase region in severing proteins does not follow a conserved pathway. Adding to these findings, in our previous studies of microtubule severing AAA+ nanomachines the hexamer in its spiral state showed a propensity for dissociation into trimers or dimers, especially in the absence of binding partners.^22,33^ In the present study, we addressed a number of questions regarding the mechanism of oligomerization/disassembly of severing proteins: (1) which oligomeric species is likely to be stable depending on conditions (ligands bound or just the APO state)?; (2) what drives the disassembly of some oligomeric species, and what keeps other oligomers intact?; (3) which secondary structural elements in the severing monomers behave as hot spots that respond to nucleotide, CTT, and/or monomer-monomer binding?; and (4) is the loss of quaternary assembly in higher order oligomers accompanied by any structural changes inside the monomers?

Our results show that the variety of essential dynamics motions supports the relative stability of oligomers, in accord with the observations based on the structural flexibility of oligomers extracted from simulations. Furthermore, the motions in the principal component space indicate that the NBD-NBD contacts are the first to be destabilized at the convex interface of a monomer, with the NBD-HBD contacts breaking later. Thus, our results show that the NBD-HBD interface between neighboring monomers is more important than the NBD to NBD interface for holding monomers together in oligomers. We also found that the NBD-NBD interface is strengthened through ligand binding. Moreover, analysis of the potential allosteric networks in severing proteins, carried out based on hot spots analysis, revealed that the HBD tip and the C-terminal helix (C-Hlx) are the common elements between the allosteric networks responding to all types of binding: nucleotide, substrate, and monomer to monomer. The clustering analysis using our newly developed StELa approach, indicated that the katanin HBD tip adopts a larger variety of secondary structures compared to the spastin tip, providing further support to the larger overall flexibility of the katanin monomer. Finally, StELa revealed that the structural ordering of the HBD tip is a characteristic of stable oligomers in both types of severing enzymes.

## METHODS

### Molecular Dynamics Simulations

#### Starting Structures

We started from the spiral and ring hexameric structures for katanin (spiral: 6UGD and ring: 6UGE) and spastin (spiral: 6P07 and ring: 6PEN) obtained in the presence of nucleotides and a minimal substrate peptide (Table S1 from the Supplementary Information).^11,12,14,34^ We split these hexameric structures into different oligomers, as depicted in Figure 1. Namely, we ran simulations for trimers, dimers, and monomers. Since the loss of nucleotide in the ring conformation occurs in monomer A for katanin and monomer F for spastin, we chose to run specific trimers and dimers for the two severing enzymes in order to probe both typical and extreme oligomers. ABC and AB are the extreme oligomers for katanin, while BC is typical for katanin. Extreme oligomers for spastin are DEF and EF. Furthermore, for each system we considered the fully complex state, where both the minimal substrate (polyE in 6UGD, 6UGE, 6P07, and polyEY in 6PEN) and ATP or ADP were bound and the APO state where the nucleotide and the substrate were removed from the original structure, following the set-ups of our previous work.^33^ Both katanin and spastin monomer simulations also included the substrate and nucleotide states, which allowed us to study the effect of removing one binding partner at a time. Each monomer contains an NBD and a HBD shown in Figure 1. These domains form crescent shapes that bend at a hinge indicated by the orange beads at residues PRO234, found in the NBD near the ATP binding site, and LEU361 which is found at the base of the HBD for katanin. Respective hinge sites for spastin are PRO525/LEU650 for 6P07 and PRO384/LEU508 for 6PEN.^11,13,14,34^ We identify the tip of the HBD, a region determined to be critical in our analysis, spans residues ASP417 to LEU437 for katanin, LEU706 to ALA721 for spastin spiral, and LEU564 to ARG581 for spastin ring.

**Figure 1:**
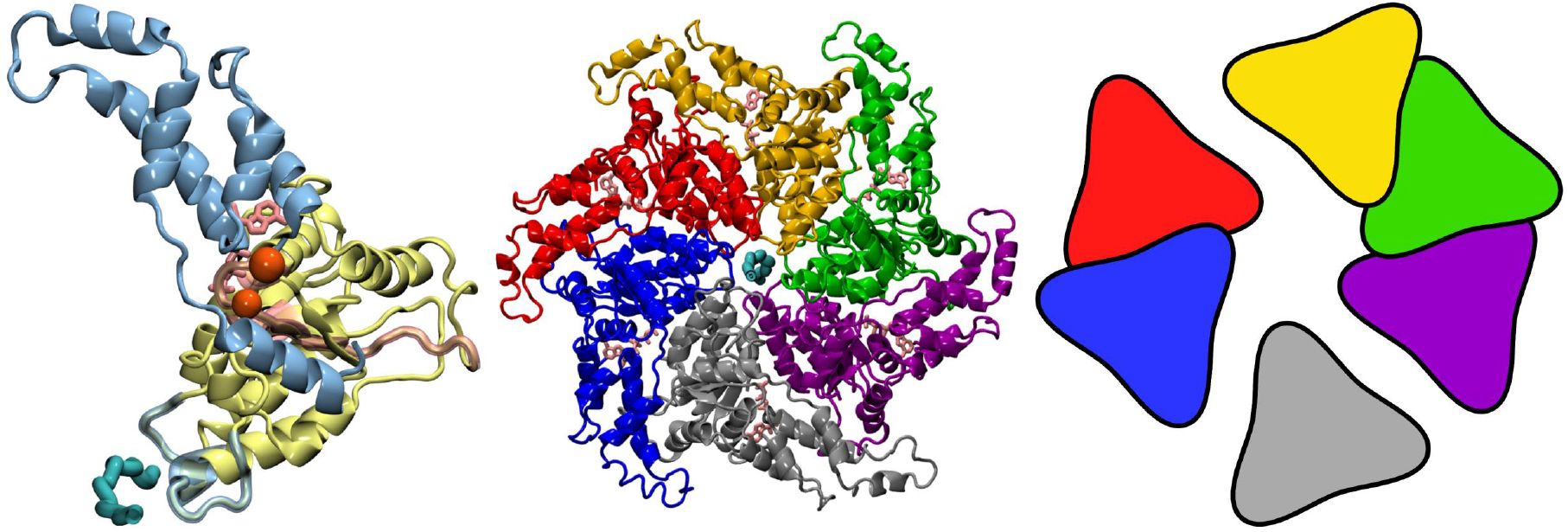
Structural details of the katanin monomer and hexamer. (left) The katanin monomer structure shows the NBD in pale yellow, the HBD in pale blue, the ATP and the ATP-binding domains in pink, and the minimal substrate and substrate binding domains in cyan. Hinge residues PRO234 and LEU361 are highlighted as beads in orange.^13^ (middle) Spiral conformation of the katanin hexamer (PDB: 6UGD)^12^ is colored by chain: monomer A (purple), monomer B (green), monomer C (gold), monomer D (red), monomer E (blue), and monomer F (grey). The ligands are colored as in the monomer on the left. (right) Cartoon representation of how we divided the hexameric assembly to extract the lower order oligomers of interest. In the katanin simulations, monomer A was extracted for the monomeric simulations, monomers AB and BC were used for the dimeric simulations, and monomers ABC were used for the trimeric simulations. In spastin simulations, monomer A was extracted for the monomeric simulations, monomers EF were used for the dimeric simulations, and monomers DEF were used for the trimeric simulations.

#### Simulation Details

We carried out full atomistic and explicit solvent molecular dynamics (MD) simulations using the GROMACS^35^ molecular modeling program version 2019 and the GROMOS96 54a7 force field.^36^ The automated topology builder server was used to generate force field parameters for ATP and ADP based on the GROMOS96 54a7 force field. ^37^ Each system was placed in the center of a cubic box (katanin ABC trimer for example: 14.67991 *nm*^3^), solvated with the single point charge (SPC) explicit solvent model, and neutralized with NaCl ions. Periodic boundary conditions (PBC) were applied in three-dimensions. The number of atoms for each system is listed in Table S2. The systems underwent energy minimization using the steepest descent algorithm and the Verlet cutoff-scheme for 50,000 steps with a criteria of the maximum force value smaller than 23.9006 kcal/mol/Å (1,000 kJ/mol/nm) to remove steric clashes. The systems were then equilibrated with the NVT ensemble to bring the temperature to 300 K using the leap-frog integrator algorithm for 500 ps. Then with the NPT ensemble to keep the system at a pressure of 1.0 bar, again with the leap-frog integrator algorithm, for 500 ps using the Parrinello-Rahman pressure coupling scheme.^38,39^ We ran four production trajectories for a minimum of 50 ns each, as detailed in Table S2. The bond lengths involving hydrogen atoms were restrained by the LINCS method to use a 2 fs integration step. A distance cutoff of 10.0 Å was applied to compute the nonbonded interactions and the Particle Mesh Ewald (PME) algorithm was used to compute the electrostatic interactions.^33^ For both spastin and katanin, we used the first protomer (A) of the hexamer for the monomeric simulations. It should be noted that, in katanin, this monomer is the monomer that lacks nucleotide density in the ring conformation.^12,14^ This is not the case in the spastin ring structure.^34^ Trajectories were visualized with VMD.^40^

### MD Analysis

#### Root-Mean-Square Deviation and Fluctuation (RMSD and RMSF)

We calculated the root mean-square distance (RMSD) of each trajectory with respect to the starting conformation using the protein backbone (see Figure S1 for an example of analysis for simulations of trimers) in order to (1) determine the convergence of our trajectories and (2) assess the overall stability of each oligomeric species. The RMSD values for each system are reported in Tables S3 and S4. We also calculated the root-mean-squared fluctuation (RMSF) from Eq. 1 for each residue and averaged the RMSF values for all trajectories per setup according to Eq. 2 to determine the flexibility of a particular region in each system relative to a reference (average) state. Figures S2 and S3 show the RMSF for katanin and spastin trimers.

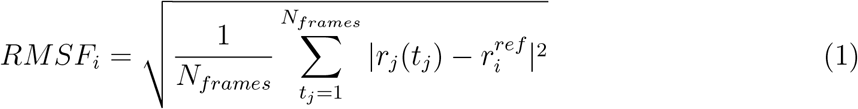

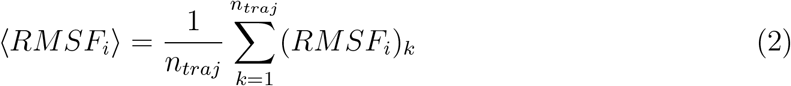

### Convergence Tests with Dynamic Cross Correlation Matrices (DCCM)

To further assess convergence of our trajectories, for each system we calculated the Dynamic CrossCorrelation Matrix (DCCM), which depicts the normalized covariance matrix and determines time-based correlations between residue pairs according to Eq. 3:

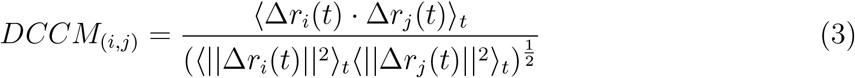

Here Δ*r*_*i*_(*t*) and Δ*r*_*j*_(*t*) are the positions of the C-*α* atoms in residues *i* and *j*, respectively, at time t. *r*_*i*_(*t*) = *r*_*i*_(*t*)− *< r*_*i*_(*t*) *>*_*t*_, *r*_*j*_(*t*) = *r*_*j*_(*t*)− *< r*_*j*_(*t*) *>*_*t*_ and *<* · *>* is the time ensemble average. Example maps for select oligomeric systems are shown in Figures S4 to S8. The convergence of the matrix was determined by evaluating Eq. 4. Here, N is the number of residue pairs and *τ* = 5*ns*.

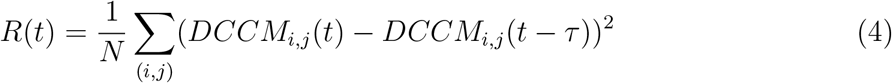

As an example, the convergence test results along with the RMSD results from katanin trimer are shown in Figure S1. These results were used to determine if the averaged trajectories were at equilibrium and therefore adequately sampled the conformational landscape. The simulations were found to have converged within ∼ 50 ns.

### Principal Component Analysis (PCA)

In order to determine the main types of motions responsible for higher RMSD values found in our simulations, we used principal component analysis (PCA) to study and compare global motions within various-sized oligomers. PCA for our systems was done by diagonalizing the covariance matrix of concatenated trajectories based on the C*α* position for each amino acid using GROMACS (gmx covar).^41,42^ This approach allowed us to reduce the dimensions of our trajectory matrix and extract the most significant modes of motion of a given system.^43,44^ We focused on only the first three components, as they represent the majority of the variance in our systems (see Tables S5 and S6 for details). We then calculated the DCCM maps for each of the three PC spaces to help discern how the motions are correlated between monomers and within monomers (see Figures S4 to S9).

### Hot Spots Analysis

We searched for secondary structures of importance in each principal component that were perturbed upon nucleotide, substrate, or monomer to monomer binding. As detailed above, the DCCM provides a map of how the motions of residues in a given system correlate with each other, and therefore offers insight into the identity of the allosteric networks of the system (see Figures S4 to S9). These maps revealed regions of high correlation (|*Corr*_*ij*_| *>* 0.9) and regions of no correlation (|*Corr*_*ij*_| *<* 0.25).^45,46^ By comparing the system without binding partners (APO) to the system with binding partners (complex), we can identify regions that become correlated (ON) or uncorrelated (OFF) upon binding, or regions that change from positively correlated to negatively correlated (EX). We identified the regions of interest for our analysis as regions in the proximity (within 12 Å) of the ATP binding site, of the Pore Loops 1 and 2, which are required for substrate binding, and of the regions at the concave and convex interfaces for a given monomer, including the HBD tip. We determined the frequency of occurrence for each residue in each list and then we combined it with the frequencies of the other residues identified for each secondary structure. In order to appropriately represent each secondary structure, we averaged the combined frequencies over the total number of residues in the structure. Finally, we identified secondary structures as significant hot spots for a given PC if they averaged above 0.500 in the monomer, and 0.300 in the dimer and trimer.

### Clustering Secondary Structures in a Selected Protein Region

Our previous analysis consistently points towards the selected HBD regions as having the most significant motions in the PC space, as well as forming interfaces important for nucleotide-binding, substrate-binding, and oligomerization. Thus, we set out to identify any potential structural differences in this region between oligomers that could further distinguish between preferred and non-preferred oligomeric states. Because the HBD tip appears often as a hot spot and both spastin and katanin monomers in the starting hexameric states show diversity within the secondary structure of this region, we focused our analysis on probing changes in the secondary structure of the HBD tip. We developed a clustering based method called StELa (<u>S</u>econdary s<u>T</u>ructural <u>E</u>nsembles with machine <u>L</u>e<u>A</u>rning) to cluster the structures of the HBD tip sampled along the various types of trajectories (see Figure S10 for method outline). The analysis starts from the Phi/Psi backbone angles of this region in each monomer.^47,48^ Next, we obtained the Ramachandran plot corresponding to these dihedral angles and we extracted the clusters for the Phi/Psi angles using the K-Means centroid based algorithm. This identified regions in the Ramachandran plot that characterize known geometries of secondary structures.^49,50^ Next, we converted each frame in a trajectory into a representative vector where the 3D atomic positions of a residue in the HBD tip are replaced by a single integer, which is the cluster index that corresponds to the secondary structure of the residue previously obtained. We then checked these representative vectors for known secondary structure propensities, such as the fact that an *α*-helix cannot be shorter than 4 consecutive residues.^51^ We used agglomerative hierarchical clustering, with complete linkage and Euclidean distance metrics applied to the extracted representative vectors, to identify the unique structures from the conformational space. The number of clusters was chosen from local maximum scores by applying three statistical methods: the silhouette score, the Calinski-Harabasz score, and the Davis-Bouldin index. Operationally, we found that the first two scores were the most relevant for our analysis. We collected the populations and representative structures determined from the representative vectors and we compared the ensembles to determine the influence of oligomerization and ligand binding on the structure of the HBD tip.

## RESULTS AND DISCUSSION

### Structural flexibility in oligomeric assemblies is highly dependent on the type of severing protein

We used the time series of the average RMSD for the various types of simulations conducted here and the simulations for the hexamers from our previous work ^33^ to compare the flexibility of the oligomeric species in katanin and spastin. Using the differences in size between oligomers and the known increase of the RMSD with the size of the protein,^52^ for each system (katanin or spastin; APO or complex; spiral or ring) we ranked the global average RMSD for each oligomeric species (monomer, dimers, trimers, hexamers) in order from highest to lowest. From this analysis we extracted the order(s) of the oligomer(s) that show an inverse trend in their global RMSD compared to the corresponding monomeric RMSD. Namely, we selected the oligomer(s) for which the RMSD is lower (or comparable) than the one for the monomer. Alternatively, if none of the higher-order oligomers average RMSD values lies below the respective monomer, we selected the oligomer with the lowest average RMSD. Finding such behavior is indicative of the stability of the respective oligomeric species.

For katanin, in the spiral conformation, there is little difference in the average RMSD between the complex and APO states in the various oligomeric assemblies (see Figure S1 and Table S3). Our simulations show that the most stable oligomer, in both the complex and APO states, was the trimer. This result is based on the fact that the average RMSD values for the trimers were in the same range as for the monomer, even when the size of the trimer is three times that of the monomer.^52^ At the other extreme, the least stable oligomer was the hexamer.^33^ In the ring conformation, for the complex state, katanin is likely found as a combination of hexamers and BC dimers. Interestingly, in the APO state, with the ABC trimer and the AB dimer being unstable, the BC dimer is the only stable oligomeric species for the katanin ring. Thus, for both the complex and APO states, the BC dimer was the most stable lower order oligomer. These findings suggest that the perturbation to the ring conformation that is associated with the absence of the nucleotide in monomer A causes significant disorder in lower-order oligomers and that the trimeric assembly that lacks a stabilizing convex interface in monomer A readily dissociates into a dimer and a monomer. For spastin, in the spiral conformation, we found that the complex was more stable compared to the APO state, which we attributed to the influence of the binding partners (Table S4). The most stable oligomer in the spiral conformation is the trimeric assembly, in both the complex and APO states, in accord with the above findings for katanin. Notably, the average RMSD of the trimers was less than the average RMSD of the monomers in each state, in spite of its much larger size, and this difference was more pronounced than in the case of katanin. The least stable oligomer was the hexameric ensemble. ^33^ In the ring conformation, where the nucleotide density was missing in monomer F,^34^ the complex and APO states have comparable average RMSD values in each oligomeric assembly. The most stable oligomer is the hexamer in both the complex and APO states. However, we note that, if the hexameric state is perturbed, the DEF trimer is the most stable lower-order oligomer in both the complex and APO states. Importantly, this behavior is very different from the one reported above for the katanin ring, where dimers are the stable oligomeric assemblies.

### Conformational dynamics of the isolated monomers reveals that katanin is intrinsically more flexible than spastin

We characterized the essential modes of domain motions for severing proteins in the monomeric, dimeric, and trimeric states using PCA. We focused on only the first three principal components, as they cover approximately 50-87% of the variance of a given system (see Tables S5 and S6). Furthermore, using the DCCM maps projected on these three modes, we determined residue pairs or regions of an assembly that undergo correlated directional motions. We found that this mode decomposition is a powerful tool for comparing the influence of the binding partners and adding interfaces to each oligomer state.

For the katanin monomer, both the APO and the substrate only states show a common directional global motion in PC1 corresponding to highly correlated large amplitude motions in the HBD (see Figure S10): a large swing-like rotation of the HBD about the hinge of the monomer (see Figure 1), which opens the crescent shape of the monomer. This correlated intradomain motion is in fact the signature behavior of the katanin monomer. Moreover, as seen in the PCs for the higher-order oligomers, such large movements in the HBD are characteristics of a monomer without a concave interface. We also found that, in PC1, the binding of the nucleotide reduces the correlations between the motions represented in the DCCMs (see Figure S4). The corresponding porcupine plot shows small amplitude local motions within each domain, signaling that the domains do not execute rigid body motions in the presence of the nucleotide (see Figure S11). This behavior is reversed dramatically upon the binding of both the nucleotide and the substrate, which induces correlations throughout the monomer resulting in highly tandem motions of the two domains. The motions of the domains as independent rigid-body units appear only in PC3 of the APO state, where the highly-correlated directional motion in the NBD is anticorrelated with the highly-correlated swing-like motion of the HBD. This mode changes dramatically upon nucleotide binding, either alone or accompanied by the substrate, to a state where the two domains move as rigid bodies and in sync with each other. We note that the DCCMs for PC2 in the presence of ligands do not change compared to the APO state, which indicates that the motions described by this principal component are not important for binding of co-factors to the monomer.

In the spastin monomer, the domain that has consistently strong directional correlations in the majority of the modes is the NBD (see Figure S5). For example, the NBD positions are highly correlated in the DCCMs for PC1 of the APO and substrate states and PC3 for the nucleotide state. The prominence of the NBD is thus a signature for the spastin monomer, which will have implications in the higher order oligomers. Like katanin, the binding of the nucleotide to the monomer introduces disorder in the PC1 DCCM, seen as the prevalence of lower amplitude local motions within and between the domains. The motions become much more correlated upon addition of the substrate, again recalling the behavior of the katanin monomer (see Figure S12). And, just like in katanin, PC3 in the presence of the nucleotide corresponds to the monomer moving as one big domain. Analysis of the directional motion using porcupine plots (see Figure 2, Figure 3 and Figures S11 to S15) shows that in each system there is also correlated movement in the HBD, as seen in the PCs for higher order oligomers. This movement is more modest than that in the terminal katanin monomer. Unlike in katanin, PC2 motions turned out to be important for substrate binding to the spastin monomer as the DCCM for PC2 in the presence of the substrate changes compared to the APO state. In fact, the binding of substrate changes the motion described by both PC1 and PC2, resulting in anti-correlated rigid body motions of the two domains. Our analysis showed that, for the katanin monomer, the binding of both the nucleotide and the substrate leads to large highly-correlated motions throughout the monomer. In contrast, the spastin monomer requires only the binding of the nucleotide to exhibit such behavior. These results signal that the monomeric katanin is intrinsically more flexible than spastin.

**Figure 2:**
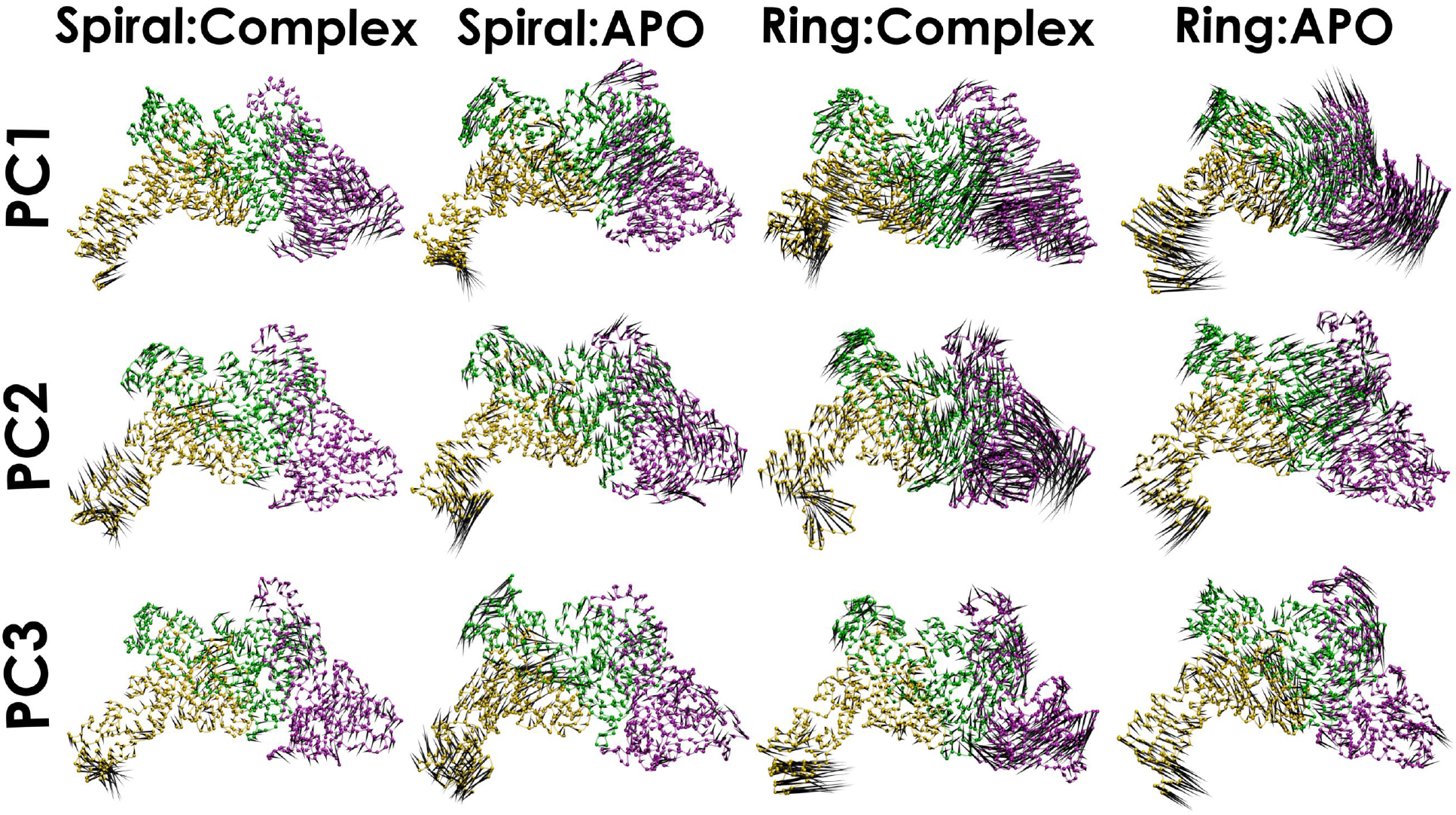
Principal component porcupine plots for the katanin ABC trimer. Colors follow the convention from Figure 1: monomer A (purple), monomer B (green), and monomer C (yellow). The PC motions are described by the black arrows with length scaled in the appropriate vector for visualization. The complex and APO setups are shown for both the spiral and ring conformations.

**Figure 3:**
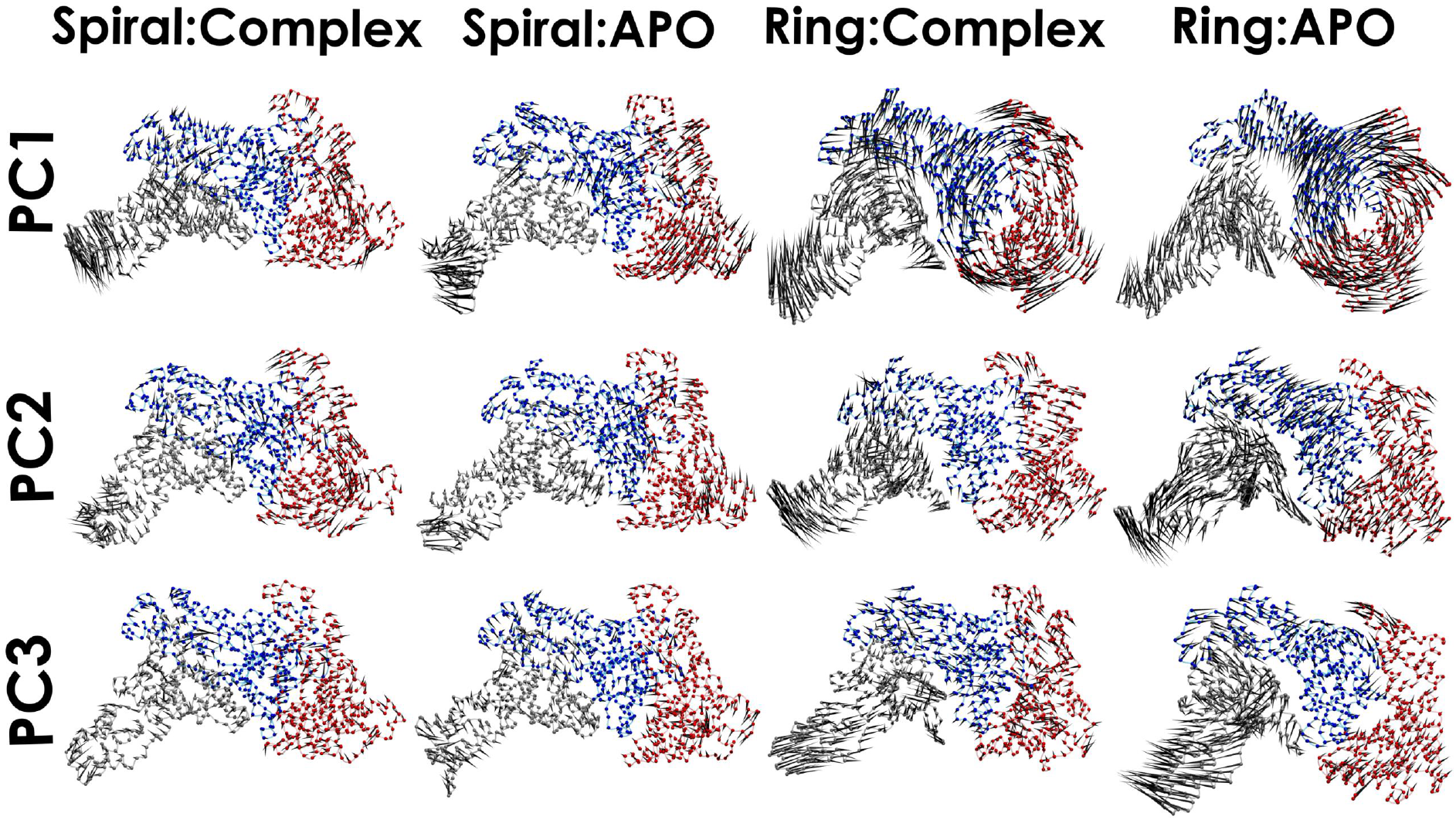
Principal component porcupine plots for the spastin DEF trimer. Colors correspond to Figure 1: monomer D (red), monomer E (blue), and monomer F (grey). The PC motions are described by the black arrows with length scaled in the appropriate vector for visualization. The complex and APO setups are shown for both the spiral and ring conformation.

### Conformational dynamics of oligomeric assemblies is highly dependent on the stability of the NBD to HBD inter-monomeric interface

For the spiral conformation of the katanin trimer in the complex state, which is stable according to the RMSD analysis, we found that the monomers execute rotational motions in PC1 leading to the rearrangement of the inter-monomer interfaces to create a more compact assembly (see Figure 2). The result is that the trimer folds in like a lawn chair due to the HBD tip of monomer C folding in about its hinge towards the A and B monomers and the NBD of monomer A rotating about its hinge towards the B and C monomers. Importantly, the highly correlated HBD motion, seen in the monomer case, is present here as well but only in monomers B and C, which are the monomers with convex interfaces. In PC2 and PC3, the motions are small rotations or rearrangements in a monomer. In the absence of both binding partners, i.e., in the APO state of the spiral conformation, PC1 shows similar compressive motions between the monomers where the HBD of monomer A moves towards the NBD of monomer B. Importantly, in both PC2 and PC3 monomer B, which is the only protomer with both neighbors present, moves like a rigid body. This recovers the behavior of the katanin monomer in the PC1 and PC3 of the complex state, detailed above. Because we did not find a similar behavior for monomer B in the spiral complex state of the trimer, we conclude that the rigidification of a katanin monomer can be achieved either due to the formation of the interfaces with both its neighboring monomers (in a trimer) or due to the binding of the nucleotide and the substrate, but not when binding all four co-factors. This conclusion is supported further by the results from the AB and BC dimers analysis, which show that the monomer with a neighbor only on its convex side, with or without the nucleotide and substrate bound, does not execute rigid-body motions.

In the motion of the complex state of the spiral AB dimer, shown in Figure S13, the HBD of monomer A moves away from monomer B in both PC1 and PC3. In the APO state, the NBD of monomer A moves away from the NBD of monomer B in PC1, while the HBD of monomer A moves away from monomer B in PC2. This separation between domains is further evidence of the instability of the dimers in the spiral conformation detected based on the RMSD analysis.

For the ring conformation of the katanin trimer in the complex state, which is unstable, we found that monomer A, which lacks ATP, rotates dramatically in PC1 such that its HBD tip moves away from the NBD of monomer B (see Figure 2). The corresponding DCCM shows that monomer A is moving anti-correlated with respect to the B and C monomers, which move as one (see Figure S8). Mode 2 shows a rotation in monomer A at the hinge resulting in part of its NBD moving away from monomer B and its HBD lifting up from the NBD of monomer B. These motions are indicative of the dissociation of monomer A from the BC dimer. Turning to PC1 of the APO state, the NBD of monomer A executes a similar large pivot rotation as in the complex state leading to the loss of NBD-NBD contacts between A and B. PC3 shows the HBD tip of monomer A peeling away from the NBD of monomer B. The losses of both the NBD-NBD and HBD-NBD interfaces would thus lead, on longer timescales, to the dissociation of the ring trimer into lower-order oligomers in both the complex and APO states. Similarly, in the complex state of the AB dimer, the motions represented in Figure S13 show that monomer A separates from monomer B in PC1. In the APO state, monomer A rotates in the same way as in the ABC trimer causing the HBD of monomer A to move away from the NBD of monomer B. These motions explain the instability in the AB dimer. In contrast, the BC dimer in the ring state was stable. In the PC1 of the complex state (Figure S14) the NBD of monomer B is a distinct region in the DCCM, while the HBD of monomer B is correlated with the motions of the NBD of monomer C. Notably, unlike the AB dimer case, the monomers do not separate in any of the PCs. A similar behavior is seen in the APO state, in support of the proposed stability of the BC dimer.

While the RMSD analysis showed a preference for katanin to be a stable trimer in the spiral conformation and at equilibrium between a dimer and a monomer in the ring conformation, spastin was stable as a trimer in the spiral conformation and at equilibrium between a hexamer and a trimer in the ring conformation. The motions corresponding to the first three PCs show a similar distinction between the severing enzymes in each oligomeric state. Whereas the katanin ring trimer displayed the loss of NBD-NBD contacts between monomers A and B in PC1 and the loss of HBD-NBD interfaces in monomers A and B in PC3, respectively, almost all spastin trimer motions preserve the monomer-monomer interfaces. Instead, in spastin we found more collaborative motions between the HBD-NBD and NBD-NBD regions of partnering monomers (see Figure 3). PC1 in the spiral conformation shows large swing-like motions in the HBD of monomer F, while PC1 in the ring conformation reflects significant global motions in the whole trimer. These large motions in the ring showed both domains in monomer D rotating counterclockwise along with the NBD of monomer E. This is similar to the PC motions of the more stable BC dimer of katanin (see Figure S14). We also found that the HBD of monomer E moves together with the NBD of monomer F. PC2 and PC3 for both spiral and ring correspond primarily to large movements within monomer F. In the complex state, monomer F’s NBD in the ring conformation moves slightly away from the NBD in monomer E, but overall the system’s PC motions reflect its stability since the HBD-NBD interface of monomers E and F stays intact. Overall, the PC motions show that the stability of the spastin trimers, inferred from their low RMSD values, can be attributed to the direction of motion of each i-th monomer following the same path as it’s (i+1)-neighboring monomer and the persistence of the HBD-NBD interfaces. The first three PCs of the spiral EF dimer, which had a higher average RMSD than the DEF trimer, show a similar loss of partnering interfaces as seen in the unstable AB dimer of the katanin ring (see Figures S13 and S15). In the ring conformation, where this dimer has the highest RMSD value compared to all other oligomers, the HBD of monomer E clearly separates from the NBD of monomer F in PC2 of the complex state and in PC3 of the APO state. Monomer F’s motion in both cases is in the opposite direction of it’s motion in the DEF trimer, which is the reflection of the absence of monomer D in the dimer. This separation also agrees with the motions found in the less-stable katanin oligomers and further supports our finding of the oligomerization’s dependence on the concave HBD and convex NBD interfaces.

### Hot Spots analysis reveals the HBD tip as the common element of allosteric networks responding to multiple co-factors binding to severing proteins

Allosteric networks mediate receptor-induced changes in conformations of proteins. Such networks usually span large distances and include inter-domain and inter-monomer interactions.^45^ As described in the Methods section, to determine the likely allosteric networks in the various oligomeric assemblies of the severing proteins, we relied on the differential entries in the DCCMs corresponding to the APO and complex states. Furthermore, to characterize long-range communication between regions, we computed the cross-correlation maps of residue fluctuations along the first three PC modes for the various systems. In many of the DCCMs from the oligomeric states we found strong correlation between motions of distant regions (see Figures S4 to S9), which supports the existence of long range interactions and coordinated domains or even full monomer movements. The structures identified to have the largest average number of hot spots upon adding various binding partners to katanin assemblies assemblies are shown in Figure 4. Each structure was numbered according the appearance in sequence as seen in Figure S16. In detail, the most significant secondary structures with respect to the nucleotide binding region, the substrate binding region, and the oligomer interfaces binding regions were identified for each principal component. By comparing these states we can determine the influence of adding binding partners to the system.

**Figure 4:**
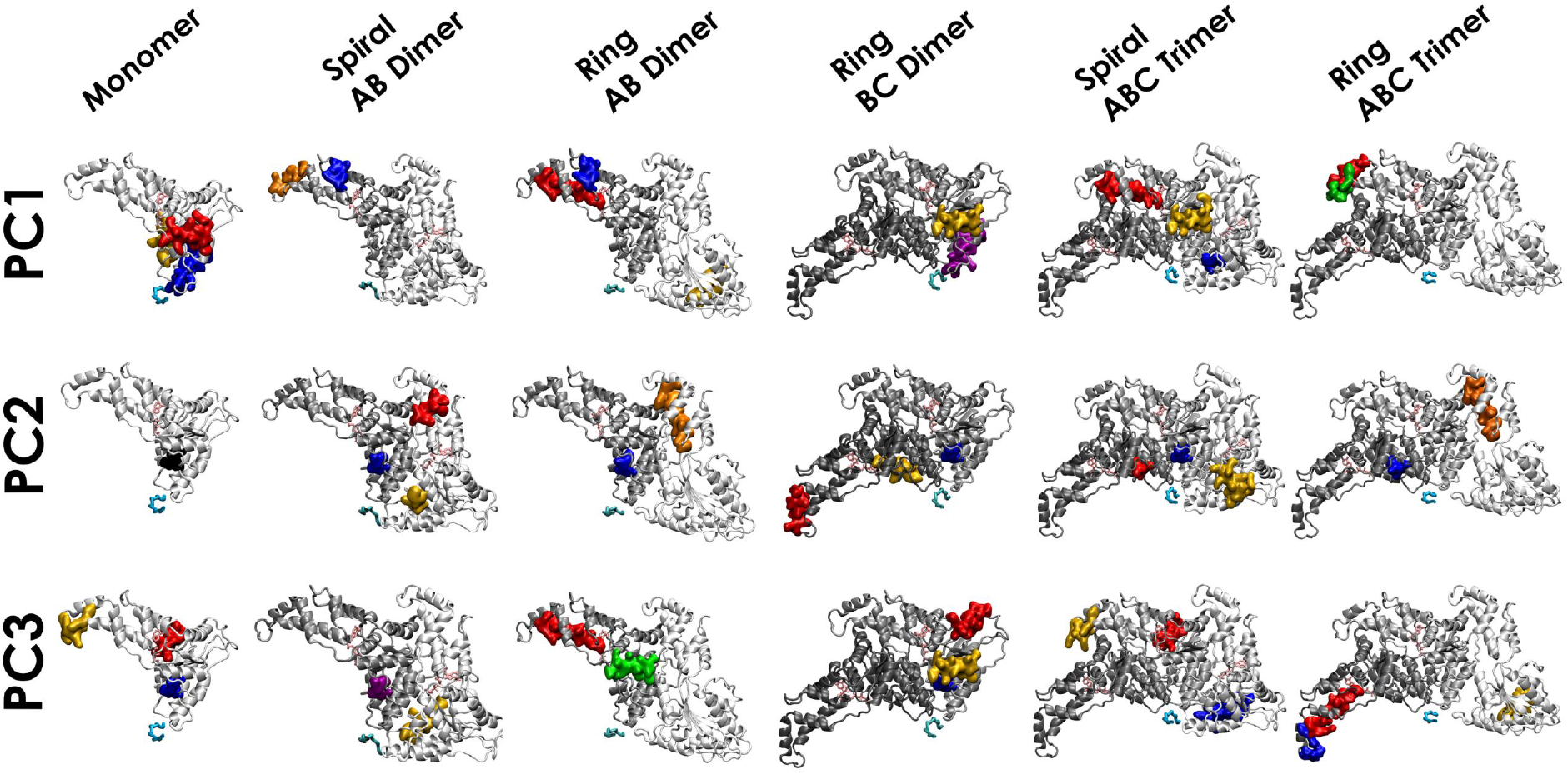
Summary of the hot spots for all the katanin oligomers. In red is the region found to be the most significant for ATP binding. In blue we highlight the region found to be the most significant for the substrate binding region. In yellow we highlight the region found to be the most significant for the formation of potential interfaces in the monomer and the existing interfaces in the dimers and trimers assemblies. Areas shared by the ATP binding region and the interface binding region are in orange. Areas shared by the substrate binding region and the interface binding regions are in green. Areas shared by the ATP binding region and the substrate binding region are in purple. The ATP molecule is highlighted in pink and the minimal substrate is highlighted in cyan.

For mode 1 in the katanin monomer we identified three independent hot spot secondary structures. The first is *α*14 (the C-Hlx), which contains the largest average number of residues that react to ATP binding. This helix wraps across the top of the monomer and packs with the Walker B region,^12,14^ which is responsible for binding and assisting in the hydrolysis of ATP. Walker B is the key region targeted when preparing the cryo-EM structures of katanin because the hexameric assembly was found to be too unstable without a point mutation (E293Q) in this region.^12,14^ The effects of this mutation are not fully understood, but using the DynaMut server we found that mutating the glutamine (Q) back to an glutamate (E) had a different effect in monomers A and E that are observed to have slightly different secondary structures in the cryo-EM (see Figure S17). ^12,53^ The predicted change in the Gibbs Free Energy from the glutamine mutation to the wild type E was -0.545 kcal/mol in monomer A, which is considered destabilizing, and +0.191 kcal/mol in monomer E, which is considered to be stabilizing (Table S7). Interestingly, this analysis showed that residues in the NBD, which would be involved in the formation of the convex interface with an adjacent monomer, and residues preceding the HBD tip, which would form a concave interface with another adjacent monomer, experienced the most dramatic deformation energy changes (see Figures S18 to S20). The second structure identified in our analysis is *α*6, which was a significant hot spot upon the binding of the substrate. This helix is located in the region right after PL2, which interacts with the substrate, and forms part of the convex interface of a monomer in the hexameric assembly.^12,14^ Finally, *α*1 had a significant number of hot spots with respect to protein-protein binding. For example, this helix is known to interact with the surface of the microtubule.^14^

For mode 1 in the spiral katanin trimer, which is the stable oligomeric species, *α*11 in monomer B was a significant hot spot upon ATP binding (see Figure 4). This was also the case in the AB spiral dimer, but not in the monomer, signaling that *α*11 becomes a hot spot reacting to ATP binding only upon the formation of the convex interface of a monomer. *α*5, which precedes PL2, was a significant hot spot in monomer A due to the substrate binding. This helix is close in sequence to the Walker B region that binds ATP, and to PL2 that binds the substrate. Its importance is also stressed by the fact that a mutation in the comparable helix of spastin is associated with hereditary spastic paraplegia.^14,18,54^ Also, our analysis shows that PL2 is more important to the allosteric network with respect to the substrate binding than PL1. The C-Hlx (*α*14) was a significant hot spot in the trimer and in the BC dimer with respect to the formation of the inter-monomer interfaces. This helix was also an important hot spot in the monomer, but with respect to the binding of ATP. Thus, we conclude that the allosteric network responding to the formation of inter-monomer interfaces in katanin oligomers has stronger connectivity than the network responding to the binding of ATP to the monomer as the latter network is easily perturbed by the formation of the oligomeric assembly. In the ring trimer, which is unstable, the HBD tip of monomer B - including the loop and the following helix - was the most significant hot spot with respect to all three types of binding regions. The only other instance of this hot spot in mode 1 is found in another unstable assembly, the AB dimer. In conclusion, hot spots in mode 1 populate the HBD in the more unstable assemblies, indicating that the HBD plays an important role in communicating with the inter-monomer interfaces.

In PC2, for all katanin assemblies, we observed a remarkable phenomenon: the same region, *α*5, is highlighted in at least one monomer with respect to the substrate binding region and, in some instances, with respect to the nucleotide binding region. This is the same helix identified in PC1 for monomer A in the katanin trimer with respect to substrate binding. The other important hot spot in PC2 is *α*11, this time only in monomer A, whose NBD is flexible. *α*11 is a hot spot with respect to nucleotide and interface binding, same as in the ring AB dimer. This helix makes up a large portion of the HBD of monomer A’s contribution to the interface with the convex face of the NBD in monomer B. As discussed above in the PC motions, the HBD to NBD interface is a crucial element whose stability can compensate for the loss of NBD-NBD contacts in order to keep monomers A and B together in an oligomeric assembly. The hot spots identified in mode 3 were regions associated with their functional roles. For example, in the isolated monomer, the helix *α*3 was significant with respect to the ATP binding region, *α*5 was significant with respect to the substrate binding region, and the HBD tip was significant with respect to the formation of inter-monomer interfaces. In monomer C of the ring trimer, *α*11 is highlighted once again with respect to the ATP binding region and *α*12 is highlighted with respect to the substrate binding region. This is the first instance that connects the HBD tip to the substrate binding region. Thus, the HBD tip is a hot spot with respect to all three types of binding addressed in our study. The only other instance of a hot spot identified with respect to all three types of binding was the C-Hlx (*α*14).

Next, we analyzed the spiral and ring conformations of katanin upon the formation of the AB and the BC dimers to determine the effect of adding either the monomer B’s convex or concave interfaces (in the AB and the BC dimers, respectively). Our results show that, compared to the isolated monomer case, upon the formation of the convex interface, the monomer (B) switches hot spots from the NBD to the HBD region. Namely, for mode 1, we found as hot spots the helices in monomer B that are close to the HBD-tip (*α*11 and *α*12). On the other hand, as mentioned above, in PC2 we found that *α*5, which includes the Walker-B motif and runs up to the start of PL2, was the primary hot spot for all oligomeric states and was specifically important upon substrate binding in all dimer set-ups. Finally, in accord with the motions depicted in the porcupine plots, for PC3 hot spots, in the spiral conformation of the AB dimer we found the Walker-B to PL2 region in the NBD of monomer B as an important hot spot due to the binding partners, while in the ring conformation the HBD-regions (*α*11 and *α*14) in monomer B become important due to inter-monomer binding. Turning to the allosteric effects due to the formation of the concave interface of a katanin monomer (in the BC dimer), we found far more modest effects compared to the formation of the convex interface. For example, in PC1 for the BC dimer we found that monomer B behaves similarly to the isolated monomer case.

Regions of spastin identified as critical for nucleotide, substrate, and interface binding had varying trends compared to katanin. Still, similar to katanin we found that the HBD tip is an important hot spot for oligomerization (see Figure 5). All three modes for the spastin monomer showcased almost identical regions: helices in the HBD region near the nucleotidebinding pocket, upon ATP binding, the long helix making up the convex interface and part of PL2, upon CTT-binding, and the HBD tip, upon interface-binding.

**Figure 5:**
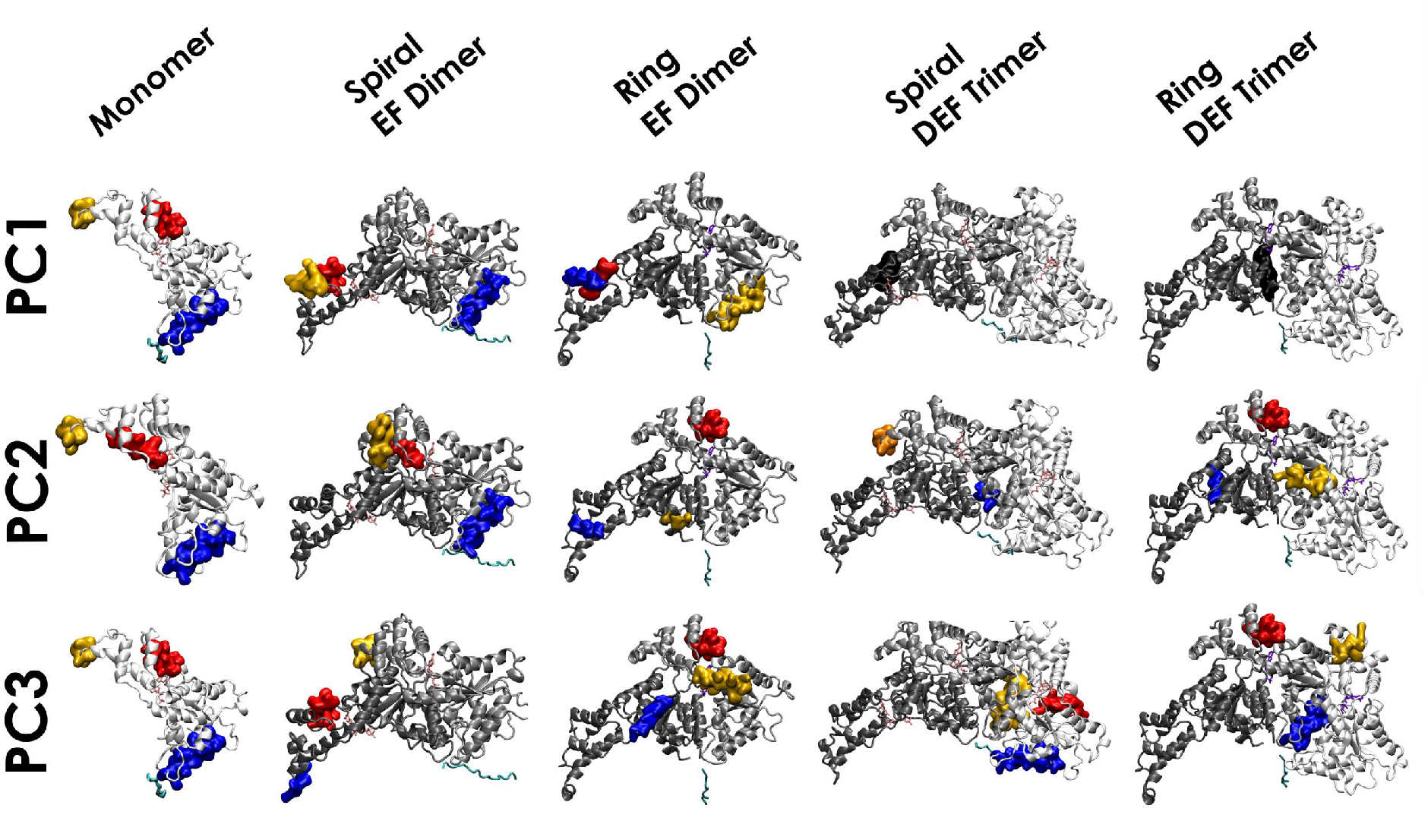
Summary of hotspots results for all spastin oligomeric states, similar to Figure 4. The red region was the most significant to the ATP (ADP if in ring conformation) binding. The blue region was the most significant to the substrate binding. The yellow region was the most significant to potential interfaces in the monomer and existing interfaces in the dimer and trimer. Regions shared by the ATP/ADP binding region and the interface binding region are indicated in orange. Regions shared by the substrate binding region and the interface binding regions are indicated in green. Regions shared by the ATP/ADP binding region and the substrate binding region are indicated in purple. Regions shared by all three binding regions are indicated in black. The ATP molecule is highlighted in pink, the ADP molecule is highlighted in violet, and the minimal substrate is highlighted in cyan.

Focusing on the EF dimer in its spiral and ring states allowed us to determine not only the effects of adding the monomer E’s concave interface, but also the difference in species, as the spastin spiral is from D.*melanogaster* and the ring from H.*sapiens* (see Methods). Interestingly, in both species we found the same hot spots regions in mode 1: HBD helices in monomer F and the PL2 helix in monomer E. The differences between species become apparent in PC2: the spiral conformation has hot spots only in monomer E, that all mimic the monomeric state, while the ring conformation places a stronger emphasis on monomer F, such as on the helix following the Walker-B motif, which becomes a hot spot upon intermonomer binding. Furthermore, in PC3 for the spiral conformation, both monomers E and F behave like the isolated monomer, while for the ring only monomer E has the same hot spots as the isolated monomer. Overall, the hot spots in the two EF dimers are consistent with the locations from the isolated monomer, irrespective of the species. This follows the trend katanin, where adding the concave interface to monomer B in the BC dimer did not cause a shift in the hot spot regions from the monomeric state. Thus, for both severing enzymes, the addition of the concave interface to a monomer does not induce any sizable allosteric effect in the monomers.

The largest allosteric signal due to the formation of inter-monomer interfaces is in the spastin trimers (Figure 5). In view of the modest role of the concave interface discussed above, we surmised that the formation of the convex interface induces allostery in the spastin systems. In detail, Figure 5 shows a clear contrast between the regions identified in the monomer/dimer structures and the trimer structures for all three PCs. Both spiral and ring conformations in PC1 showed that the binding of any of the co-factors resulted in only one hot spot in monomer F’s NBD region. In the spiral, the hot spot was a small helix in the NBD of monomer F, which wraps itself on the top of the ATP-binding pocket, and does not appear in any other oligomer (*α*2). In the ring conformation the hot spot was a part of PL2 in the NBD, which was also critical for CTT binding in the monomeric state. In PC2, monomer E in the spiral conformation behaved similarly to the spiral dimer by highlighting the HBD tip, while identifying the helix in PL1 as important for CTT-binding. The ring also showed monomer E as having one region similar to the ring dimer and the rest unique. PC3 for the spiral conformation is the only state to reveal an important hot spot in monomer D, which was the PL2 region identified in the monomer.

### Clustering analysis of the oligomeric conformations shows that the HBD tip readily undergoes secondary structure changes

The hot spots analysis for both severing proteins identified the HBD tip as the most prevalent region in allosteric networks connected with the binding of all the various co-factors, as detailed above. In addition, we found that the HBD tip undergoes dramatic fluctuations (see Figures S2 and S3). Importantly, our analysis showed that the stability of any oligomeric species in severing proteins is highly dependent on the persistence of the HBD-NBD inter-monomer interface, which involves the HBD tip. When these contacts are lost, the oligomers dissociate. These results strongly indicate that the HBD tip must have important functional roles in severing enzymes, which are likely dependent on its structure. Thus, next we focused on the characterization of the secondary structure of the HBD tip in our trajectories.

Analysis of the cryo-EM structures of the hexamers (Figure S17) shows that, in both katanin conformations (6UGD and 6UGE), the HDB tip of monomers A, B, C, and F contains a two turn helix, while in monomers D and E there are only flexible loops. ^12,14^ The HBD tip of the spastin spiral (6P07) contained a short helix followed by a flexible loop in monomers A, B and E while in monomers C, D and F, there were only flexible loops. ^11^ In the spastin ring (6PEN), the comparable region is a short helix in all 6 monomers.^34^ The differences among monomers indicate that this region is characterized by a relatively low conformational energy barrier.

Next, we used PSIPRED to predict the secondary structure of the HBD tip based on the amino acid sequence.^55^ PSIPRED requires a sequence of at least 35 amino acids and the HBD tip region in each severing protein is up to 21 amino acids in length. Thus, to PSIPRED we instead provided a longer sequence, centered on the HBD tip (see Figure S21). The prediction for katanin from C.*elegans* is that a flexible loop forms after the long helix, followed by a short helix, and another short flexible loop. A nucleotide free monomer for H.*sapiens* solved by McNally et. al shows two small helices in the HBD tip region. ^13^ Using the H.*sapiens* katanin sequence, PSIPRED predicted two helices in this region instead of a single small helix, in accord with the crystal structure of the monomer. The prediction for both spastin from D.*melanogaster* and from H.*sapiens* indicates that, following the long HBD helix, there is a short helix and a flexible loop. The prediction matches the solved structures for spastin relatively well, aside from the HBD tip regions in the monomers that were solved as unstructured.

In order to better characterize the conformational variability in the HBD tip of the two main severing proteins, we developed a clustering method, StELa, based on the backbone torsion angles of the amino acids for this region extracted from our simulations using GROMACS (see Methods for details). By comparing the systems we can see the influence of the different ligand binding and interfaces on the HBD tip region. The majority of the variation in the structures for a given system was found in the length of the starting helix, as well as the angles of the loop regions. Interestingly, we identified six types of structures in the katanin assemblies (see Figure 6) and only four types of structures in the spastin assemblies (see Figure 8), which are distinguished by the number and location of individual helices. The populations of these structures were used to assess changes between oligomeric assemblies and ligand binding as seen in Figure 7 and Figures S22 to S31. The number of unique structures and representative structures with their corresponding populations identified through StELa for the katanin monomer are shown in Figures S32 and S33. StELa revealed that there are only two HBD tip conformations found in each ligand state (see Figures S22 to S27). The first was the starting structure found in monomer A, the Loop-Helix structure. ^12,14^ In the complex state, over 98% of the structures are Loop-Helix structures. In the nucleotide state about 74% of the structures resembled the Loop-Helix structure, while in the substrate state we found this structure 39% of the time. In the absence of any binding partners, 88% of the structures adopt the Loop-Helix arrangement. The second conformation was the Helix-Helix structure, seen in Figure 7. We found this conformation primarily in the presence of the substrate only (60%), or the nucleotide only (25%), while being virtually non-existent in the complex state (1%). The HBD tip was correlated with both ligand binding sites in the hot spots analysis, which probably accounts for the binding-dependent structural changes in this region. Namely, the independent binding of the nucleotide or the substrate caused the flexible loop to form a helix. When both ligands are bound to the system, this region reverts to the flexible loop found in the APO state. This result supports the finding based on the PC motions that the two ligands have canceling effects on the system. The presence of the Helix-Helix conformation was especially striking as it resembles the two helix HBD tip found in the “auto-inhibited” H.*sapiens* monomeric crystal structure.^13^ The complex state yielded two more flexible HBD tip conformations at populations less than 1%: Loop-Helix-Helix and Helix-Helix-Helix. The Loop-Helix-Helix and Helix-Helix-Helix ensembles were unexpected as they are the results of a helix breaking in half. The break (containing at least 2 non helical residues with at least 4 helical residues before and after) was predominantly observed between residues 428 and 431 (ALA-ALA-MET-GLU sequence). The Loop-Helix-Helix structure also resembles the two helices found in the auto-inhibited monomer. ^13^ This suggests there may be two different pathways to achieve the conformations of the HBD tip from the H.*sapiens* monomers. One where the loop transitions to a helix, which is the preferred pathway, and one where the helix breaks resulting in the formation of the two helices. In view of the fact that the overwhelming majority of the structures resemble the starting structure in the APO and complex state, the transition is likely due to the influence of the binding partners.

**Figure 6:**
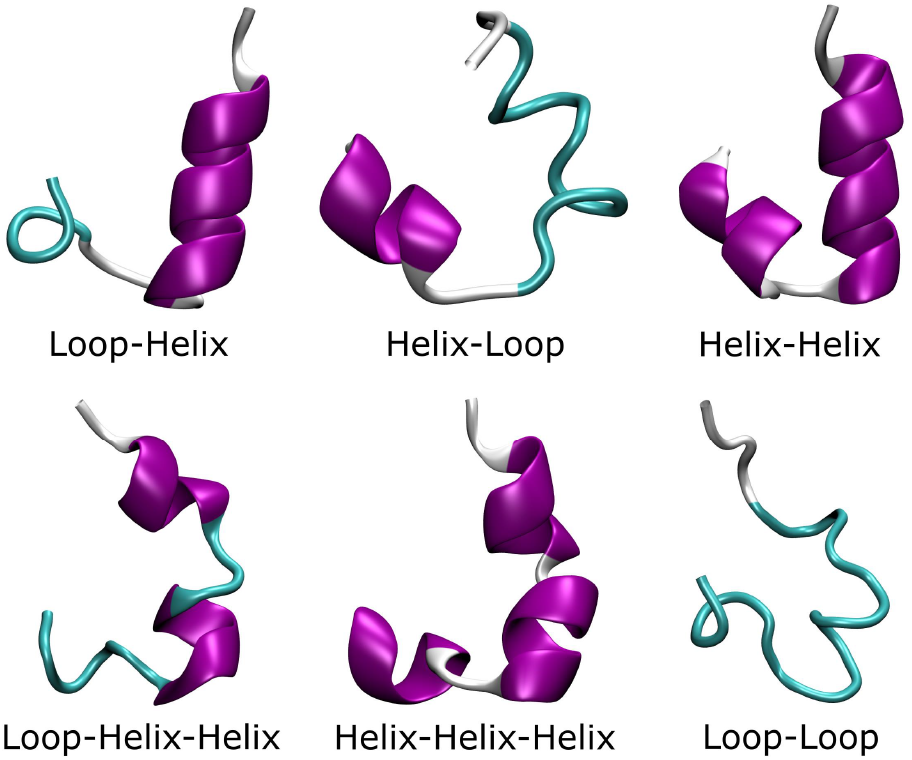
The six representative types of structures found in the simulations of katanin systems using the clustering approach StELa.

**Figure 7:**
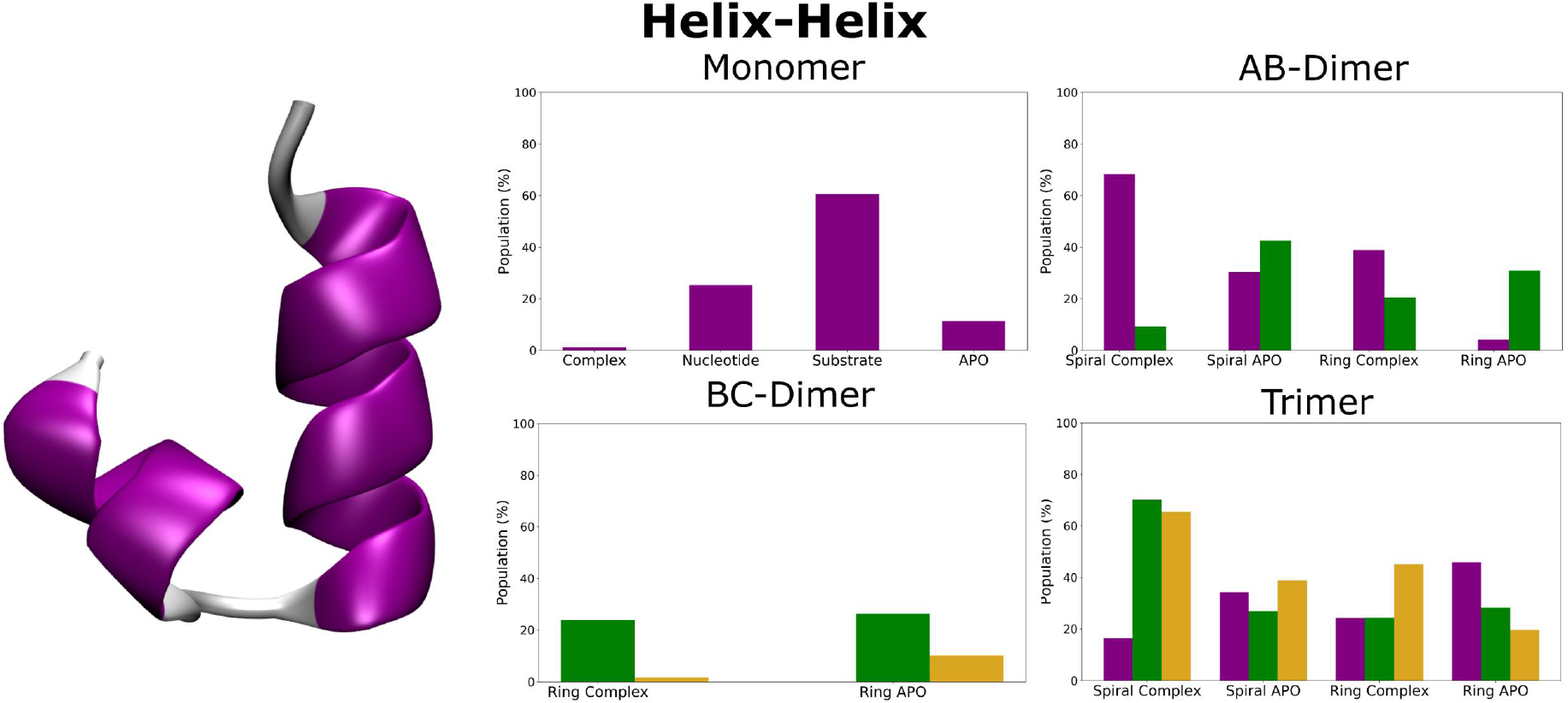
The populations found for the Helix-Helix structure of the HBD tip in the various katanin systems using StELa. The structure is shown on the left. The title of each graph indicates the type of oligomer. The purple, green, and yellow bars indicate populations for monomer A, monomer B, and monomer C, respectively.

**Figure 8:**
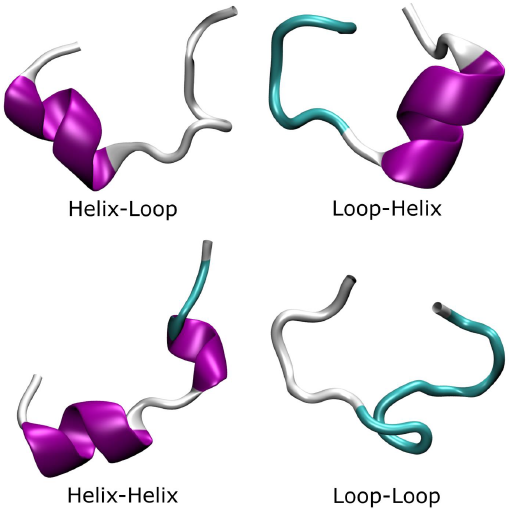
The representative structures of the four types of structures found in the simulations of spastin systems with StELa.

In the dimeric assemblies, we see some of the trends determined from the monomeric simulations, but with some differences (see Figures S22 to S27). For the AB dimer, we found an elevated population of the Helix-Helix structure in both states of the spiral conformation (67% and 32% for complex and APO, respectively) and the complex state of the ring conformation for monomer A (41%), and in both states of the ring conformation (20% and 28% for complex and APO, respectively) and the APO state of the spiral conformation in monomer

B. In the BC dimer, we identified small populations of novel varieties of more flexible structures (Helix-Loop and Loop-Loop) (see Figures S22 to S27). As these structures are not found in the monomer, we inferred that they are the result of the presence of inter-monomer interface(s). The AB dimer, which was substantially unstable in both the spiral and ring states, revealed three conformational variations in the HBD tip exclusively in monomer A in the APO state. In this assembly, monomer A forms a concave interface with monomer B where the HBD of A is bound to the NBD of monomer B. Because we know this assembly to be unstable, the resulting flexibility may be due to the relative movement of the these neighboring monomers. In the BC dimer, the Helix-Loop and Loop-Loop conformations had low populations all below 2% in the complex state of monomer B. The Loop-Loop and Loop-Helix-Helix conformations had even lower populations in the APO state of monomer B. In contrast, in monomer C, which does not have a concave interface in this assembly, had a relatively larger population of the Loop-Helix-Helix conformation in the complex state (20%).

In the spiral conformation of the ABC trimer, monomer A recapitulates the behavior of the isolated monomer where the majority of the HBD tip structures correspond to the starting structure in both the complex (83%) and APO (65%) states (see Figures S22 to S27). There was a slight elevation in the probability of the tip populating the Helix-Helix conformation in comparison to the monomeric assembly (Complex: 16% and APO: 34%). This contributes to the idea gleaned from the behavior of monomers in the dimers that the concave interface alone leads to added flexibility in the HBD tip region; however, in the trimer, the populations of the varied structures were dramatically reduced compared to the dimer case. For monomers B and C in the APO state, the HBD tip populated overwhelmingly the starting conformation (B: 73% and C: 61%). In contrast, in the complex, the HBD tip populates the Helix-Helix conformation (B: 70% and C: 65%). Because monomers B and C in the trimer have in common the presence of their convex interface, which we determined to be stabilizing for the HBD tip in the hot spots analysis, suggests that the combined effect of the formation of the convex interface and the binding of both ligands drives the HBD tip to adopt the more ordered Helix-Helix conformation. Moreover, because in the trimer we hardly see any of the more flexible structures in the HBD tip region, we propose that the adoption of the Helix-Helix conformation by this region is a signature of the stability of the trimeric assembly in the spiral conformation. This conclusion is further supported by our finding that in the ring conformation of the ABC trimer, which is unstable, the HBD tip populates the various conformations with similar probabilities as in the isolated monomer.

Next, we used StELa to analyze the region making up the HBD tip for each of the spastin oligomers. Overall, we observed four types of secondary structural conformations for spastin found in Figure 8, namely a Helix-Loop, Loop-Helix, Helix-Helix and a continuous flexible loop (Loop-Loop) (see Figures S28 to S31). These are fewer than the structures found in katanin, most likely due to spastin’s HBD tip being shorter and therefore unable to adopt the Helix-Helix-Helix and Loop-Helix-Helix ensembles observed in katanin. It is important to note that although there was less diversity found in the types of possible structures in this region for spastin, the HBD tip in katanin oligomers populated mostly only two of the six conformations (Loop-Helix and Helix-Helix), while in spastin the HBD tip populated with much higher probability each of the four identified structures, depending on state. This trend was found for all oligomers and shows that the conformation of the HBD tip in spastin does not depend on the long helix towards the C-term as it does in katanin.

For the spastin monomer, SteLa yielded that over 70% of the representative structures are in the Helix-Loop ensemble, regardless of spastin’s state. This shows a relatively high consistency in that region, regardless of binding state, since the starting cryo-EM structure shows the Helix-Loop in all monomers of the spiral (see Figure S17). In the presence of ATP, the monomer populated more structures, including Helix-Helix if the substrate was also bound (∼25%) and Loop-Loop if just ATP was present (∼20%). This is of interest since there were larger fluctuations in the monomer’s HBD tip PC motions, while the hot spots analysis repeatedly identified the HBD tip as important upon interface-binding. We observed that if the interface is important to this region and its secondary structure changes based of the inclusion of binding-partners, the HBD tip makes an important connection between ATP/CTT-binding and spastin oligomerization.

Along with observing a change in the monomer based on binding partners, when both the convex and concave interfaces are formed in the DEF trimer, there seems to be even more regularity in the secondary structure of both the spiral and ring conformations, compared to the monomer. This concurs with the very low global RMSD values found in both conformations of the trimeric state, even when compared to the monomer. The HBD tips of monomers D and E both have a Helix-Loop in their starting structure and keep this conformation throughout our simulations. The complex trimers did not show the same decrease in Helix-Loop population that the monomeric form did, further suggesting that the stability of the trimers could be due to the stability of structure the HBD tip. Monomer F starts as a Helix-Loop in the ring state, while it is unstructured in the spiral state (see Figure S17). In the ring DEF trimer, the HBD tip of monomer F has the same trend as in monomers D and E. However, in the spiral DEF trimer, the HBD tip of monomer F changes from Loop-Loop to Loop-Helix. This change from an unstructured to a more stable Loop-Helix structure, gives further support for the proposal that the increased stability of each monomer’s HBD tip is a characteristic of stable oligomeric species.

## CONCLUSIONS

Previously, we found that the hexameric form of the spiral conformation was unstable, while the ring hexamer was stable in both spastin and katanin.^33^ In the current work, analysis of our long atomistic MD simulations of spiral oligomers led us to infer that, for both systems, the trimer was the preferred stable oligomer based on the relatively lower global RMSD average in comparison to the dimer and monomer. We also found that the spastin trimer was the preferred stable oligomer in a perturbed ring conformation. In katanin, the ring trimer was substantially unstable, which was likely due to the lack of ATP in monomer A. ^14^ This indicates that the ring trimer would further dissociate into the BC dimer and the A monomer. Strikingly, this suggests that, for katanin, the process of oligomerization is distinct from the process of dissociation.^13,14,31,32^

The analysis of the PC motions provided microscopic insight into the origin of oligomers stability. The direction of the motions for the regions involved in NBD-NBD and NBDHBD connections were compressive in setups corresponding to lower RMSD values where interfaces were sealed up, whereas setups with higher RMSD values were more likely to display motions leading to the separation of interfaces. In both katanin and spastin, stable oligomers retained the HBD-NBD interactions, even when the NBD-NBD interactions were lost, while the HBD-NBD connections were lost in several unstable setups, pointing towards the crucial role played by the HBD in oligomeric stability.

Using the hot spots analysis we found that, in both katanin and spastin, the formation of the convex interface between neighboring monomers in an oligomer induces shifts in the allosteric networks. For example, secondary structure elements that were part of an allosteric network responding to the binding of the ATP in the isolated katanin monomer, switch to an allosteric network responding to monomer-to-monomer interface binding in the spiral trimer. We found that the convex and concave interfaces play different allosteric roles in the two severing enzymes. Namely, in mode 1 of the katanin spiral dimers we found that all three hot spots were located in the HBD of the second monomer (B), where in the isolated monomer all were located in the NBD. In contrast, for the spastin spiral dimers we found only two hot spots in the HBD of the second monomer (F), with the third hot spot being located in the NBD of the first monomer (E), closely resembling the behavior seen in the isolated spastin monomer. Keeping in mind that the second monomer in a dimer does not have a concave interface neighbor, while the first monomer lacks a convex interface neighbor, we infer that the presence of the convex neighbor is crucial for the allosteric networks of a monomer in spastin, while in katanin the presence of the concave neighbor is more important. This potentially gives an indication of directionality to adding protomers during oligomerization. Another interesting finding is that, in the katanin ring trimers, the hot spots were all located at the HBD tip. As this assembly was determined to be dramatically unstable based both on the RMSD comparison and the PC motions and thus doomed for dissociation, we inferred that finding all the hot spots at the HBD tip is a signature of structural instability in katanin. For spastin, where both the spiral and ring trimers were stable, we identified a single internal (i.e., not at a potential interface) hot spot in the third monomer, which possesses only a convex interface. This lends additional support to the important role played by the convex interface neighbor in the allostery of spastin’s monomers. In summary, the hot spots analysis hints to differences between the function of katanin versus spastin on microtubules.^56^ One of the most significant recurring hot spots in katanin, but not in spastin, was the CHlx (*α*14). Importantly, literature results identified this region as unique to severing enzymes from other AAA+ proteins found in the same meiotic clade. ^12–14^ This helix extends from the HBD of a given monomer and wraps across the top of its NBD, thereby creating a belt around the C-terminal side of the channel with neighboring protomers.^14^ In addition, this helix was proposed to play a significant role in oligomerization due to its packing with the Walker B region and the first helix in the sequence at the bottom of the monomer.^14^ Finally, it has been found that this helix undergoes a rearrangement from the pseudo hexameric state to the monomeric nucleotide-free state.^13^ A previously reported study of katanin sequences shows an overall 35% conservation from H.*sapiens* to C.*elegans*.^56^

In spite of the noted differences, a common finding for both katanin and spastin is that in both severing enzymes the helices associated with PL2 are more significant to the allosteric network with respect to the substrate binding than that helices associated with PL1. While both PL1 and PL2 have been determined essential for severing activity and make up a double spiral ladder around the electronegative substrate, PL1 has been established to be essential for recognizing and binding the substrate.^2,12,14,18,57^ This finding, combined with our results, implies unique roles for each of the pore loops where PL1 pulls in the substrate and PL2 is responsible for communicating the action to the rest of the motor. In the katanin hexamer, each of the pore loops are in contact with the substrate in the cryo-EM structure and in tight coordination with the formation of the interfaces between neighboring monomers implying a close relationship between the substrate binding, the ATPase activity, and the oligomerization.^12^ Our analysis shows that the substrate binding region is connected to the ATP binding regions and regions along interfaces. Finally, we found that the HBD tip is the only major hot spot associated with the binding of all three co-factors to a given monomer.

The characterization of the secondary structure of the HBD tip in our simulations proved to be an important aspect for understanding what drives the formation of the allosteric network in each severing protein. We found a strong dependence on the presence of the various ligands and provided molecular details in support of the degree of stability of the oligomeric states. In detail, using our newly developed clustering approach, StELa, we characterized states that are likely to contribute to a conformational change in the secondary structure of the HBD tip from a loop to an alpha-helix, especially in the absence of a bound nucleotide.

In addition, we observed that the conformational transition differs between katanin and spastin, adding a new layer to the idea that the two severing proteins follow distinct functional pathways. In katanin the substrate state and nucleotide state each promote the Helix-Helix ensemble, but in the complex they “cancel out”. In spastin the presence of the nucleotide alone causes a loss of structure resulting in the formation of the Loop-Loop structure, while the complex state exhibits the opposite effect where the HBD tip populated the Helix-Helix ensemble. The substrate alone had no impact on this region. Thus, in spastin, the ligands are cooperative. In the spiral trimer of katanin, which is the stable oligomer, the only structures found for the HBD tip of all three monomers were the original state (Loop-Helix) and the Helix-Helix state found in the monomer. No other structures were populated in the complex state, which was found to be the most stable. Very small populations of the Loop-Helix-Helix variety were found in monomers A and B of the APO state, likely due to the lack of ligand binding. Meanwhile in the ring trimer, which was substantially unstable and suffered a loss of key interactions between monomers, generally lower populations of the Helix-Helix ensemble were found and the more flexible structures were found with higher propensities. Interestingly, monomer B experienced the largest presence of these more flexible ensembles. This suggests that as monomer A peels away from monomer B, but monomer B stays attached to monomer C, the HBD tip of monomer B has to become more flexible in order to stay attached to monomer C.

## Acknowledgement

We thank Mangesh Damre for help with setting up the GROMACS simulations for katanin. We thank George Stan for stimulating discussions on the oligomerization mechanism in Clp proteins. This research was funded by the National Science Foundation (NSF) MCB1817948 (to RID). D.G. and S.M were supported through the NSF Research Experience for Undergraduates in Chemistry grant CHE-1950244. This work used the Extreme Science and Engineering Discovery Environment (XSEDE) through allocation TG–BIO210094 to RID.

## Supporting Information Available

The Supporting Information is available free of charge on the ACS Publications website at. Supplemental data analysis; Table S1, list of PDB information used in our simulations; Table S2, list of MD simulations and the simulation information; Figure S1, Example of convergence tests; Table S3, Global average RMSD values for each system of katanin; Table S4, Global average RMSD values for each system of spastin; Figure S2, The average RMSF plot for each residue in katanin; Figure S3, The average RMSF plot for each residue in spastin; Table S5, Percentage of variance for the first three principal components for each system of katanin; Table S6, Percentage of variance for the first three principal components for each system of spastin; Figure S4, The DCCM for each PC for the katanin monomer systems; Figure S5, The DCCM for each PC for the spastin monomer systems; Figure S6, The DCCM for each PC for the katanin AB-Dimer and BC-Dimer; Figure S7, The DCCM for each PC for the spastin EF-Dimer; Figure S8, The DCCM for each PC for the katanin ABC-Trimer; Figure S9, the DCCM for each PC for the spastin DEF-Trimer; Figure S10, the StELa clustering scheme; Figure S11, The porcupine plots for each PC of the katanin monomers; Figure S12, The porcupine plots for each PC of the spastin monomers; Figure S13, The porcupine plots for each PC of the katanin AB-Dimer; Figure S14, The porcupine plots for each PC of the katanin BC-Dimer; Figure S15, The porcupine plots for each PC of the spastin EF-Dimer; Figure S16, The secondary structure reference for the hot spots analysis; Figure S17, the cryo-EM structures for the katanin and spastin used in simulation with highlighted HBD tips; Figure S18, The DynaMut deformation energy profiles of the Walker B Q to E mutation for monomers A and E of C.*elegans* katanin; Figure S19, The DynaMut deformation energy profiles of the Walker B Q to E mutation for monomers A and C of D.*melanogaster* spastin; Figure S20, The DynaMut deformation energy profiles of the Walker B Q to E mutation for monomer B of H.*sapiens* spastin; Table S7, The outcome ΔΔ*G* predictions from the DynaMut server for the Walker B mutation; Figure S21, The PSIPRED prediction for the secondary structure of the HBD tip; Figure S22, The population of states with a Loop-Helix structure identified to be the starting HBD tip conformation in katanin; Figure S23, The population of states with a Helix-Helix structure identified in katanin HBD tip; Figure S24, The population of states with a Helix-Loop structure identified in the katanin HBD tip; Figure S25, The population of states with a Loop-Loop structure identified in the katanin HBD tip; Figure S26, The population of the states with a Loop-Helix-Helix structure identified in the katanin HBD tip; Figure S27, The population of the states with a HelixHelix-Helix structure identified in the katanin HBD tip; Figure S28, The population of states with a Helix-Loop structure identified to be the starting HBD tip conformation in katanin; Figure S29, The population of states with a Helix-Helix structure identified in the spastin HBD tip; Figure S30, The population of states with a Loop-Helix structure identified in the spastin HBD tip; Figure S31, The population of states with Loop-Loop structure identified in the spastin HBD tip; Figure S32, The representative structure for the top three populated clusters found in the katanin monomer; Figure S33, The representative structure for the top three populated clusters found in the spastin monomer.

## TOC Graphic

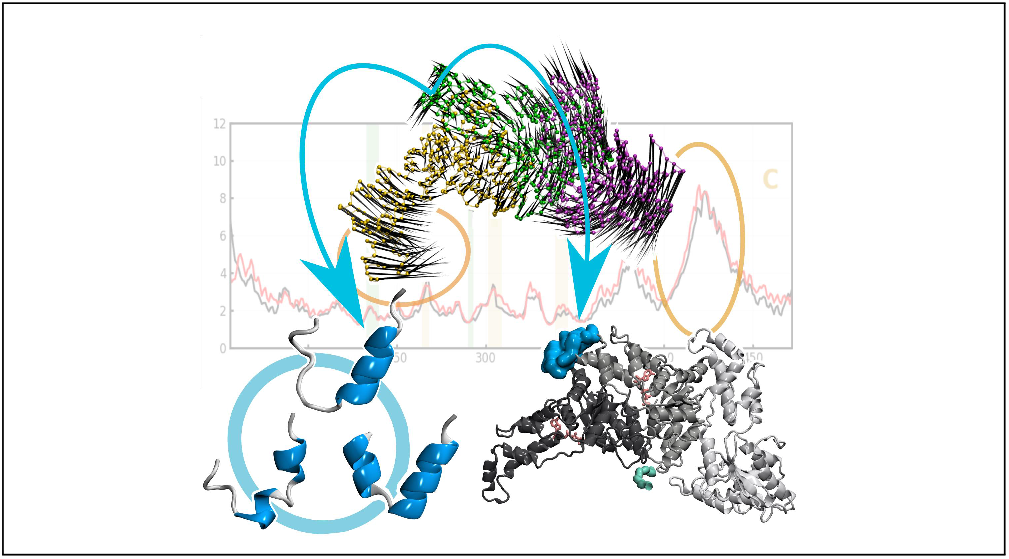

