## Supplemental tables and figures for "Microtubule severing enzymes oligomerization and allostery: a tale of two domains"

| Protein | PDB ID | Species | Resolution (A) | State | Substrate | Nucleotide |
|---|---|---|---|---|---|---|
| **Katanin** | 6UGD | *Caenorhabditis elegans* | 3.5 | Spiral | Poly-Glutamate (E14) | ATP |
| | 6UGE | *Caenorhabditis elegans* | 3.6 | Ring | Poly-Glutamate (E12) | ATP |
| **Spastin** | 6P07 | *Drosophilia melanogaster* | 3.2 | Spiral | Poly-Glutamate (E15) | ATP |
| | 6PEN | *Homo Sapiens* | 4.2 | Ring | $(EY)_5$ | ADP |

**Table S1.** PDB information for the initial katanin and spastin hexamers in the spiral and ring conformations used for all simulations.[1-4]

| Protein | Oligomer | Nucleotide | Substrate | # Atoms | # Residues | # Water (SOL) | # Na Ions | # Trajectories (Time) | Total Simulation Time |
|---|---|---|---|---|---|---|---|---|---|
| **6UGD** | Monomer | | | | | | | | |
| | A | ATP | E14 | 105031 | 331 | 33892 | 24 | 3 (50ns) | 150 ns |
| | | ATP | - | 105084 | 317 | 33962 | 10 | 3 (50ns) | 150 ns |
| | | - | E14 | 105050 | 331 | 33914 | 21 | 3 (50ns) | 150 ns |
| | | - | - | 105118 | 317 | 33989 | 7 | 3 (50ns) | 150 ns |
| | Dimer | | | | | | | | |
| | AB | ATP | E14 | 236719 | 648 | 76722 | 34 | 4 (50ns) | 200 ns |
| | | - | - | 236834 | 634 | 76844 | 14 | 3 (50ns) | 150 ns |
| | Trimer | | | | | | | | |
| | ABC | ATP | E14 | 312703 | 965 | 100984 | 44 | 4 (50ns) | 200 ns |
| | | - | - | 312849 | 951 | 101132 | 21 | 4 (75ns) | 300 ns |
| **6UGE** | Dimer | | | | | | | | |
| | AB | ATP | E12 | 286438 | 646 | 93318 | 29 | 4 (50ns) | 200 ns |
| | | - | - | 236834 | 634 | 76844 | 14 | 3 (50ns) | 150 ns |
| | BC | ATP | E12 | 248805 | 646 | 80758 | 32 | 3 (50ns) | 150 ns |
| | | - | - | 248879 | 634 | 80859 | 14 | 3 (75ns) | 225 ns |
| | Trimer | | | | | | | | |
| | ABC | ATP | E12 | 327052 | 963 | 105790 | 39 | 4 (100ns) | 400 ns |
| | | - | - | 855078 | 951 | 392986 | 21 | 4 (100ns) | 400 ns |

<table>
<tr><th>Protein</th><th>Oligomer</th><th>Nucleotide</th><th>Substrate</th><th># Atoms</th><th># Residues</th><th># Water (SOL)</th><th># Na Ions</th><th># Trajectories (Time)</th><th>Total Simulation Time</th></tr>
<tr><td rowspan="13">6P07</td><td colspan="9">Monomer</td></tr>
<tr><td rowspan="4">A</td><td>ATP</td><td>E15</td><td>102124</td><td>319</td><td>32974</td><td>22</td><td>3 (50ns)</td><td>150 ns</td></tr>
<tr><td>ATP</td><td>-</td><td>102217</td><td>304</td><td>3306</td><td>7</td><td>3 (50ns)</td><td>150 ns</td></tr>
<tr><td>-</td><td>E15</td><td>102119</td><td>319</td><td>32988</td><td>19</td><td>3 (50ns)</td><td>150 ns</td></tr>
<tr><td>-</td><td>-</td><td>114740</td><td>304</td><td>37251</td><td>4</td><td>3 (50ns)</td><td>150 ns</td></tr>
<tr><td>F</td><td>-</td><td>-</td><td>105884</td><td>304</td><td>34299</td><td>4</td><td>3 (50ns)</td><td>150 ns</td></tr>
<tr><td colspan="9">Dimer</td></tr>
<tr><td rowspan="2">EF</td><td>ATP</td><td>E15</td><td>238931</td><td>623</td><td>77565</td><td>29</td><td>3 (50ns)</td><td>150 ns</td></tr>
<tr><td>-</td><td>-</td><td>239029</td><td>608</td><td>77685</td><td>8</td><td>4 (50ns)</td><td>200 ns</td></tr>
<tr><td colspan="9">Trimer</td></tr>
<tr><td rowspan="2">DEF</td><td>ATP</td><td>E15</td><td>310392</td><td>924</td><td>100374</td><td>36</td><td>4 (50ns)</td><td>200 ns</td></tr>
<tr><td>-</td><td>-</td><td>310515</td><td>909</td><td>100518</td><td>12</td><td>4 (75ns)</td><td>300 ns</td></tr>
<tr><td rowspan="9">6PEN</td><td colspan="9">Monomer</td></tr>
<tr><td>F</td><td>-</td><td>-</td><td>115375</td><td>288</td><td>34299</td><td>4</td><td>3 (50ns)</td><td>150 ns</td></tr>
<tr><td colspan="9">Dimer</td></tr>
<tr><td rowspan="2">EF</td><td>ATP</td><td>$(EY)_5$</td><td>216684</td><td>298</td><td>70270</td><td>8</td><td>3 (50ns)</td><td>150 ns</td></tr>
<tr><td>-</td><td>-</td><td>216773</td><td>288</td><td>70363</td><td>0</td><td>3 (50ns)</td><td>150ns</td></tr>
<tr><td colspan="9">Trimer</td></tr>
<tr><td rowspan="2">DEF</td><td>ATP</td><td>$(EY)_5$</td><td>323328</td><td>876</td><td>104839</td><td>11</td><td>4 (50ns)</td><td>200 ns</td></tr>
<tr><td>-</td><td>-</td><td>349514</td><td>866</td><td>113645</td><td>0</td><td>4 (75ns)</td><td>300ns</td></tr>
</table>

**Table S2.** Summary of simulation setups, including each conformation of katanin and spastin, various sized oligomers, and the removal of one or both nucleotide and substrate.

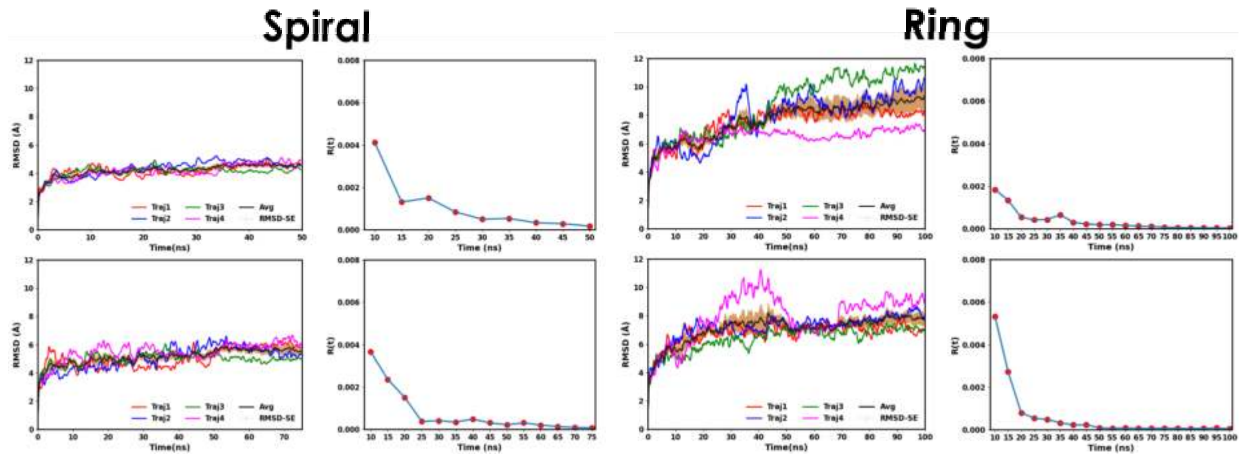


**Figure S1**. Example of the convergence tests from katanin ABC trimer simulations following the approach used in our previous studies.[5] Plots in first column show the root-mean-square deviation (RMSD) over time, with each trajectory colored independently. The average of the four trajectories is in black, and the standard error for the trajectory is shadowed in yellow. The left figures show the DCCM convergence test 5 ns apart. The system was considered converged if the last 3 points were close to 0. The example shows the test for the katanin trimers for both conformations (spiral and ring)  as well as for both the complex (top row) and APO (bottom row) states.

| **Spiral** | | | **Ring** | | |
|---|---|---|---|---|---|
| *APO* | | | | | |
| | Global Average (A) | Last 5 ns Error | | Global Average (A) | Last 5 ns Error |
| Monomer - A | 4.856 | 7.95% | Dimer - AB | 7.092 | 9.33% |
| Dimer - AB | 5.058 | 7.55% | Dimer - BC | **5.967** | 4.88% |
| Trimer - ABC | **5.131** | 4.83% | Trimer - ABC | 7.070 | 5.85% |
| Hexamer | ~6.5 | | Hexamer | ~7.5 | |
| *Complex* | | | | | |
| Monomer - A | *4.921* | 8.14% | Dimer - AB | 7.325 | 7.19% |
| Dimer - AB | 5.341 | 10.08% | Dimer - BC | **4.134** | 3.67% |
| Trimer - ABC | **4.188** | 2.57% | Trimer - ABC | 7.637 | 8.80% |
| Hexamer | ~6.5 | | Hexamer | **~5.5** | |

**Table S3.** Global average RMSD values for each oligomer of katanin spiral and ring in either the complex or the APO state. The standard error of the final 5 ns was used to determine whether simulations had properly converged. RMSD values bolded indicate the oligomers for each set up with the lowest global average. The global averages for the hexamer simulations are taken from our previous study for comparison.[5]

| **Spiral** | | | **Ring** | | |
|---|---|---|---|---|---|
| *APO* | | | | | |
| | Global Average (A) | Last 5 ns Error | | Global Average (A) | Last 5 ns Error |
| Monomer - A | 6.176 | 12.45% | Monomer - F | 5.667 | 4.27% |
| Monomer - F | 5.346 | 7.32% | Dimer - EF | 6.268 | 3.46% |
| Dimer - EF | 5.663 | 12.55% | Trimer - DEF | 6.205 | 7.94% |
| Trimer - DEF | **4.105** | 8.04% | Hexamer | **~6.0** | |
| Hexamer | ~7.0 | | | | |
| *Complex* | | | | | |
| Monomer - A | *4.309* | 6.06% | Dimer - EF | 6.264 | 9.15% |
| Dimer - EF | 3.810 | 1.75% | Trimer - DEF | 5.235 | 14.07% |
| Trimer - DEF | **3.967** | 3.26% | Hexamer | **~5.0** | |
| Hexamer | ~7.0 | | | | |

**Table S4.** Global average RMSD values for each oligomer of spastin spiral and ring in either the complex or the APO state, similar to Table S3. RMSD values bolded indicate the oligomers for each set up with the lowest global average. The global averages for the hexamer simulations are taken from our previous study for comparison.[5]

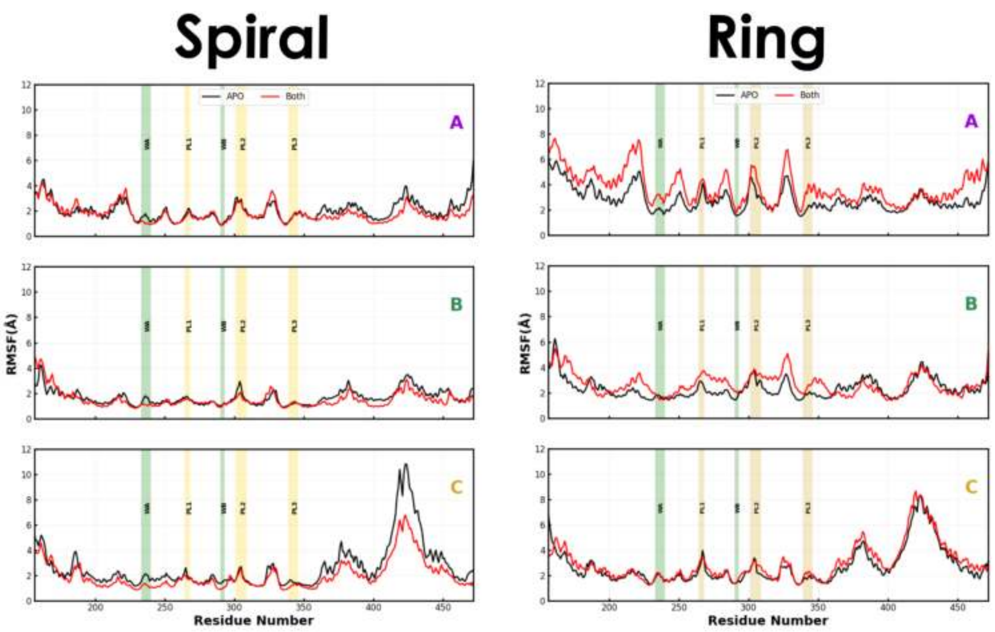


**Figure S2**. The average root-mean-sqaure fluctuation (RMSF) plot of each residue for all katanin ABC trimer simulations. The complex state RMSF is plotted in red, while the APO state is in black. The Walker-A and Walker-B motifs are highlighted in light green, and the pore loops (PL1, PL2, and PL3) are highlighted in light yellow. Each row of plots represents monomer A, B, and C, respectively, for the spiral and ring conformation of katanin.

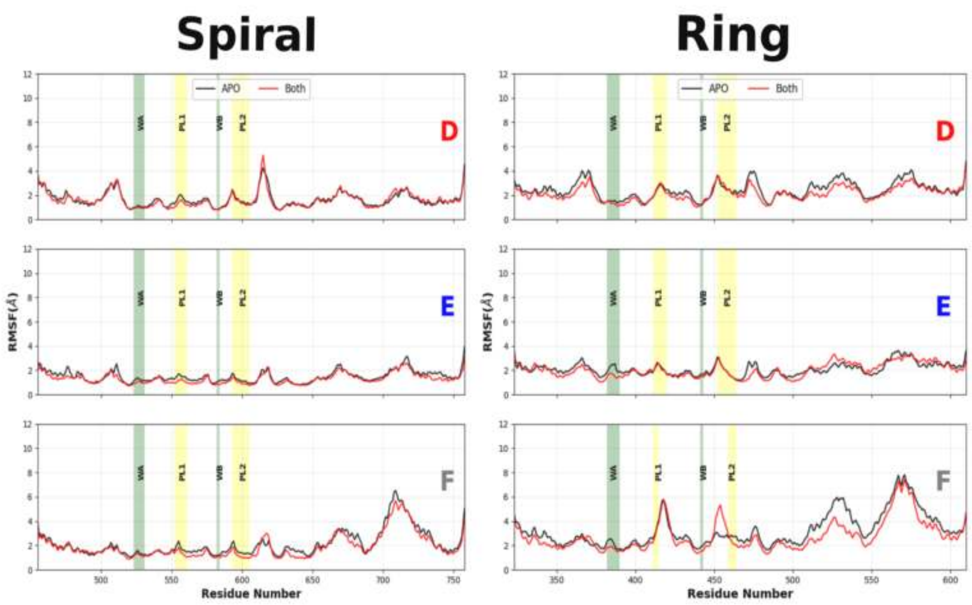


**Figure S3**. Similar RMSF plots as in Figure S2 this time for all spastin DEF trimer simulations. Each row of plots represents monomer D, E, and F, respectively, for both conformations of spastin.

| Spiral | | | | | |
|---|---|---|---|---|---|
| | | *PC1* | *PC2* | *PC3* | *SUM* |
| A | *Complex* | 42.4% | 21.8% | 7.1% | 71.3% |
| | *ATP* | 59.3% | 11.2% | 5.8% | 76.3% |
| | *Substrate* | 50.5% | 15.6% | 7.0% | 73.1% |
| | *APO* | 40.3% | 18.3% | 16.2% | 74.8% |
| AB | *Complex* | 27.4% | 12.3% | 7.4% | 47.0% |
| | *APO* | 35.3% | 13.0% | 7.9% | 56.2% |
| ABC | *Complex* | 28.3% | 24.1% | 16.1% | 68.5% |
| | *APO* | 25.9% | 21.5% | 17.6% | 65.0% |
| **Ring** | | | | | |
| | | *PC1* | *PC2* | *PC3* | *SUM* |
| AB | *Complex* | 31.0% | 14.2% | 8.3% | 57.5% |
| | *APO* | 29.0% | 17.4% | 8.5% | 55.0% |
| BC | *Complex* | 26.2% | 12.0% | 7.6% | 45.8% |
| | *APO* | 69.3% | 7.5% | 3.4% | 80.2% |
| ABC | *Complex* | 27.1% | 26.1% | 15.6% | 68.8% |
| | *APO* | 42.7% | 19.9% | 10.7% | 73.3% |

**Table S5**. Percentage of variance described by the first three principal components for all trajectories in each oligomeric setup of katanin.

| Spiral | | | | | |
|---|---|---|---|---|---|
| | | *PC1* | *PC2* | *PC3* | *SUM* |
| A | *Complex* | 43.1% | 17.4% | 7.4% | 67.9% |
| | *ATP* | 44.6% | 17.6% | 9.2% | 71.4% |
| | *Substrate* | 58.1% | 22.5% | 5.0% | 85.6% |
| | *APO* | 41.5% | 28.2% | 4.8% | 74.5% |
| F | *APO* | 58.5% | 19.9% | 3.4% | 81.8% |
| EF | *Complex* | 25.3% | 11.1% | 8.3% | 44.7% |
| | *APO* | 38.7% | 10.4% | 6.2% | 55.3% |
| DEF | *Complex* | 42.6% | 16.0% | 12.6% | 71.2% |
| | *APO* | 36.2% | 19.1% | 11.2% | 65.5% |
| **Ring** | | | | | |
| | | *PC1* | *PC2* | *PC3* | *SUM* |
| F | *APO* | 49.3% | 25.3% | 7.7% | 82.3% |
| EF | *Complex* | 50.5% | 16.3% | 6.9% | 73.7% |
| | *APO* | 37.1% | 15.0% | 7.0% | 59.0% |
| DEF | *Complex* | 40.7% | 30.8% | 15.5% | 87.0% |
| | *APO* | 36.8% | 31.4% | 11.0% | 79.2% |

**Table S6**. Percentage of variance described by the first three principal components for all trajectories in each oligomeric setup of spastin.

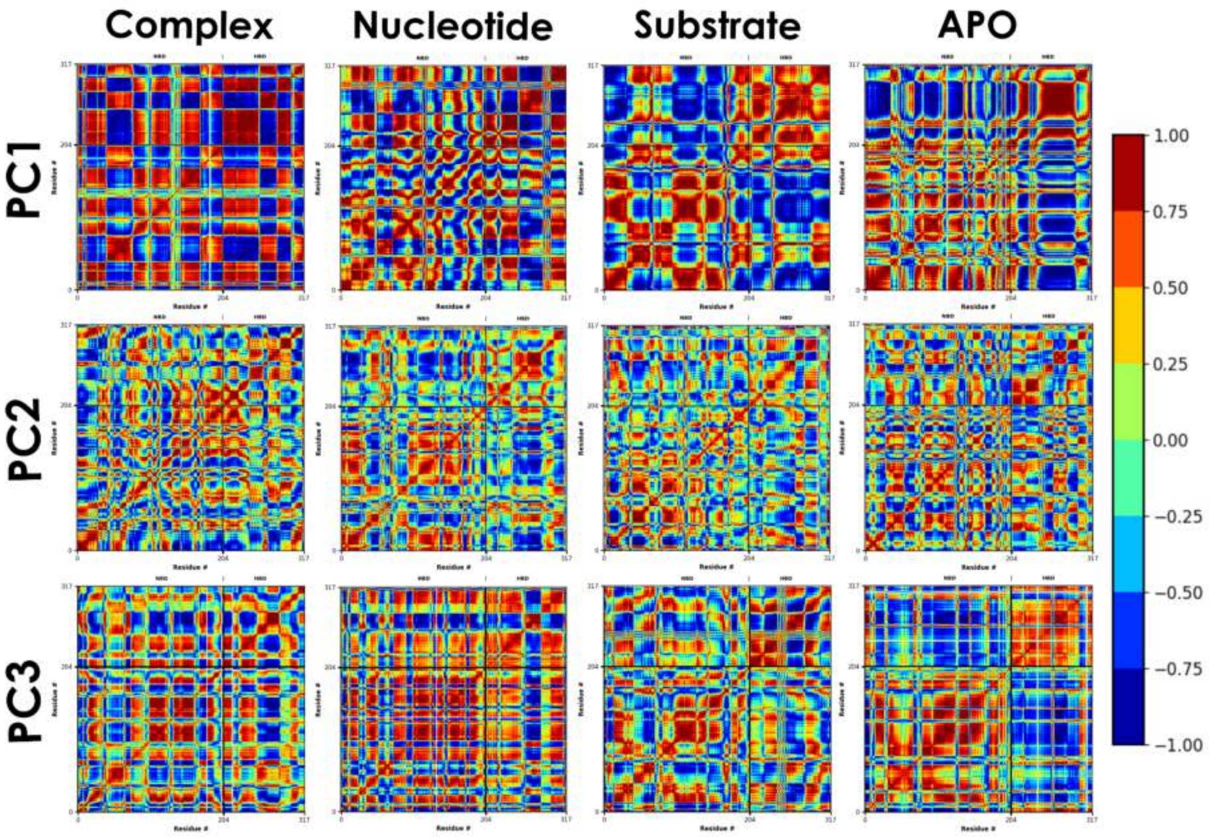


**Figure S4**. The DCCM for the first three PC for the complex, nucleotide, substrate and APO states of the katanin monomer.[6] The changes in color show how each of the ligands affects the allostery of the protein. The deep red color represents residue pairs that are highly correlated (+0.9 to +1.0) while the deep blue regions represent residue pairs that are highly anti-correlated (-0.9 to -1.0). The green regions represent residue pairs that are considered uncorrelated (+0.25 to -0.25).[7] The changes of color patterns and color blocks between the complex state to the APO state give indication of how the motion of the system is influenced by the various factors of interest. These changes are used as a signal in the hot spots analysis and demonstrate how the different influences (ligands, interfaces, and conformation) affect the allosteric network of the system.

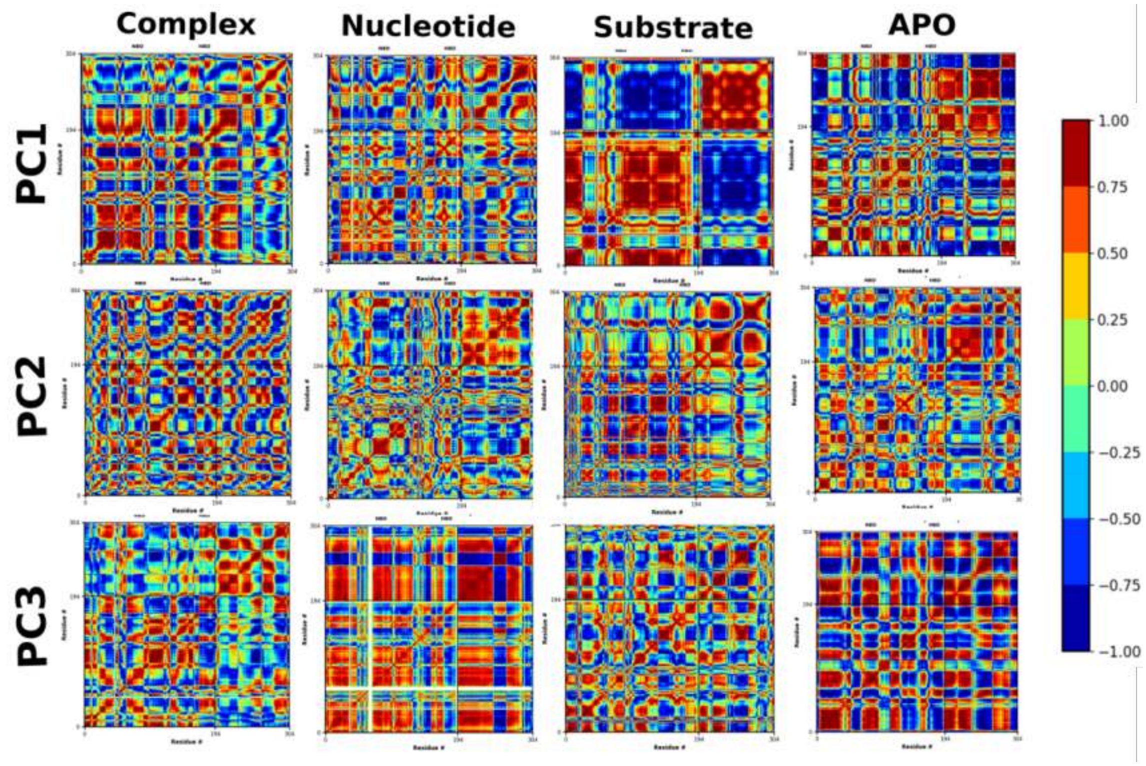

**Figure S5**. The DCCM of the first three PCs. Similar to Figure S4, but for the spastin monomer.

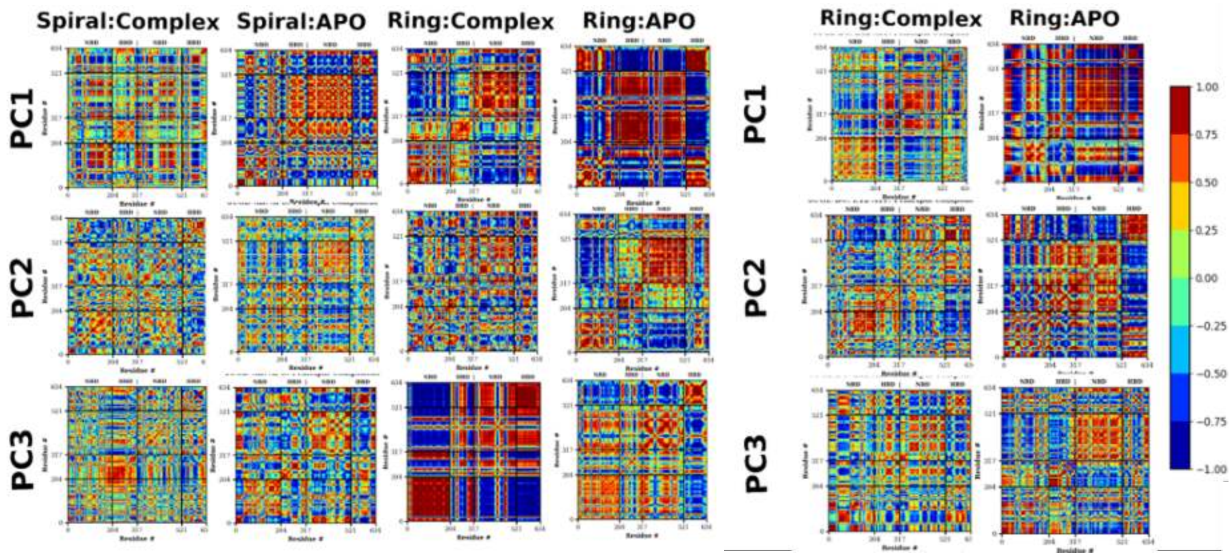


**Figure S6**. The DCCM of the first 3 PCs. Similar to Figure S4, but for the complex and APO states of the katanin spiral and ring (6UGD) AB-dimer (left) and ring (6UGE) BC-dimer (right).

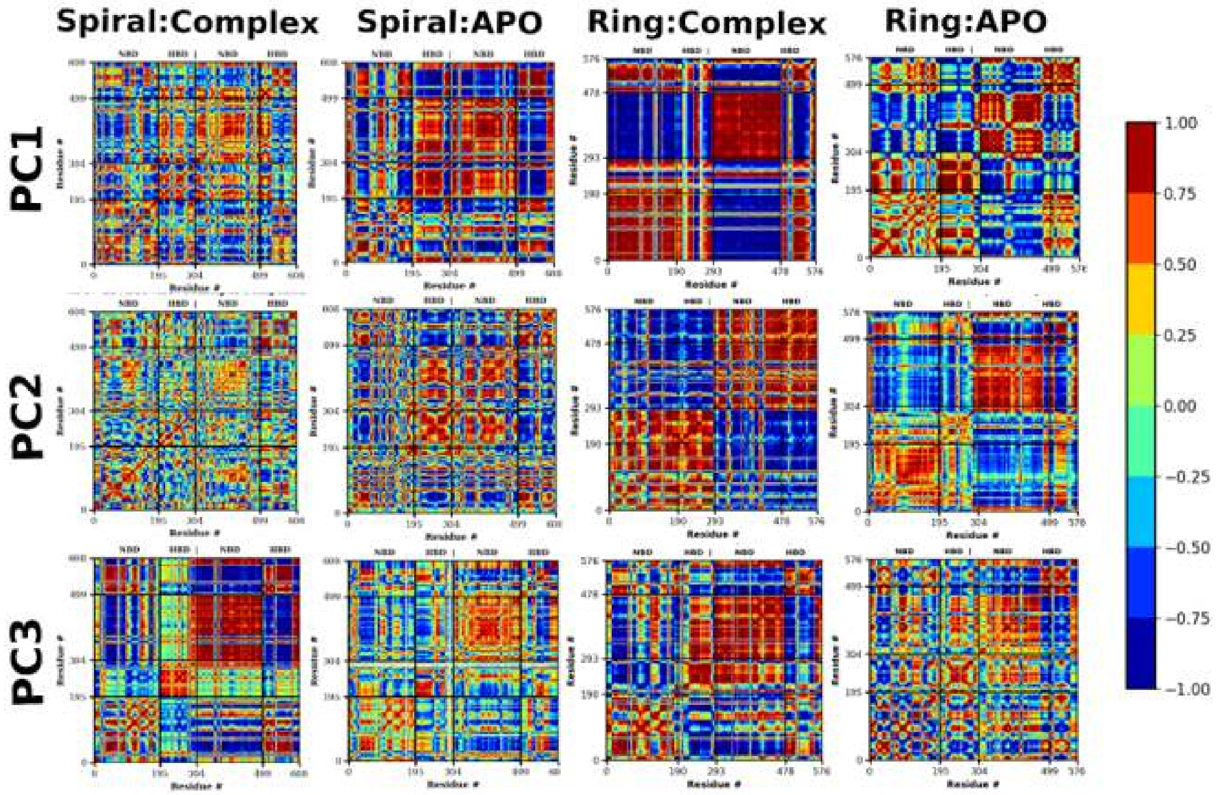


**Figure S7**. The DCCM of the first 3 PCs. Similar to Figure S4, but for the complex and APO states of the katanin spiral and ring EF-dimer.

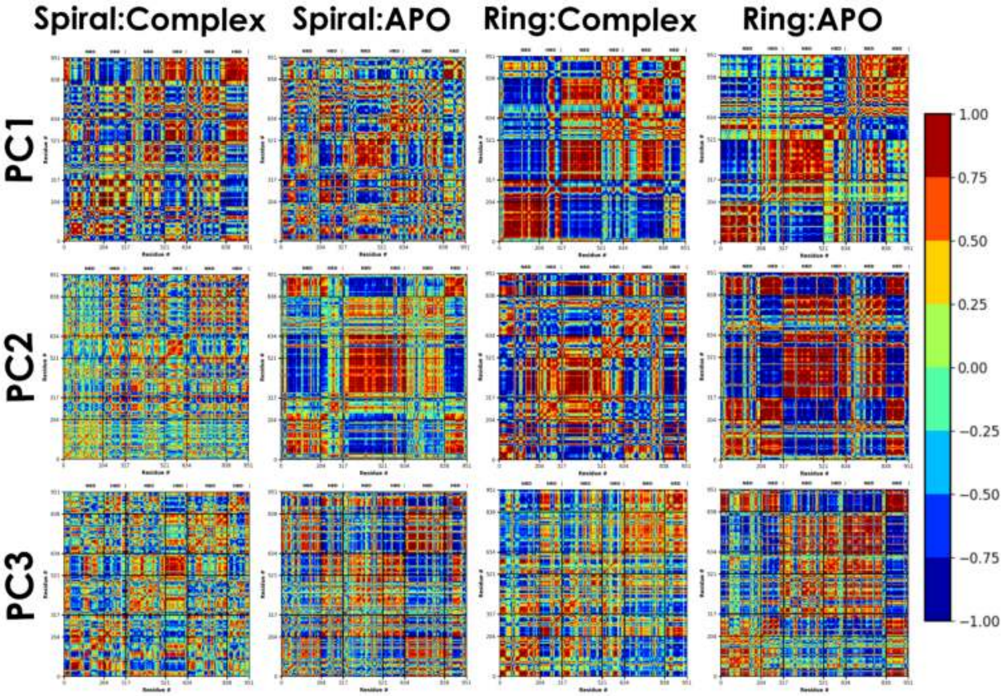


**Figure S8**. The DCCM of the first 3 PCs. Similar to Figure S4, but for the complex and APO states of the katanin spiral (6UGD) and ring (6UGE) ABC-trimer.

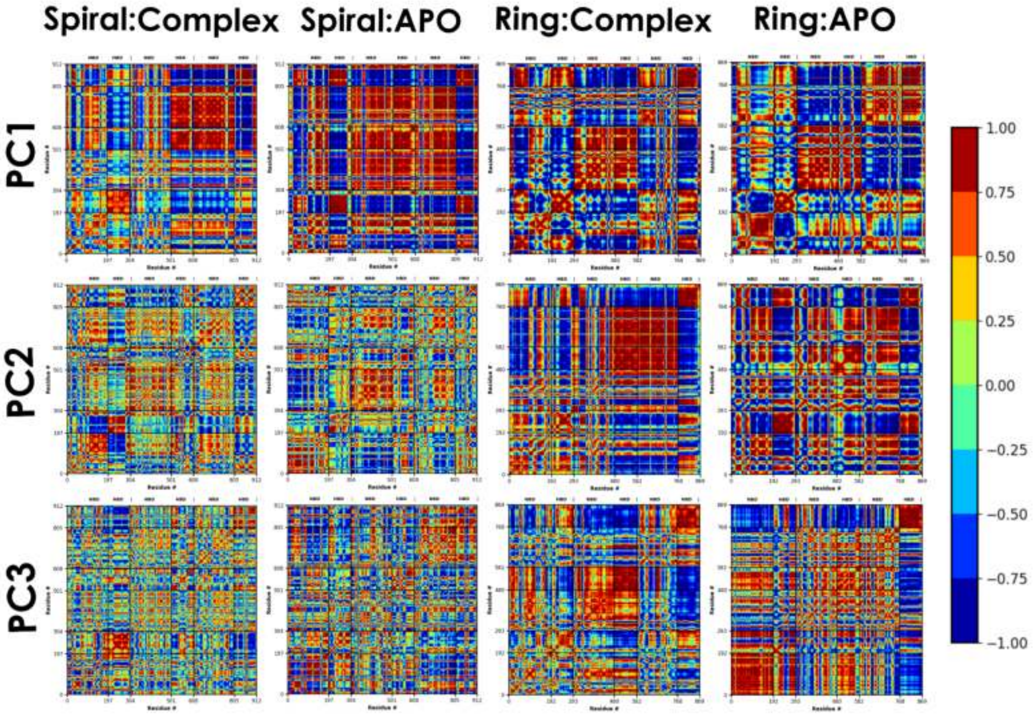


**Figure S9**. The DCCM of the first 3 PCs. Similar to Figure S4, but for the complex and APO states of the spastin spiral (6P07) and ring (6PEN) DEF-trimer.

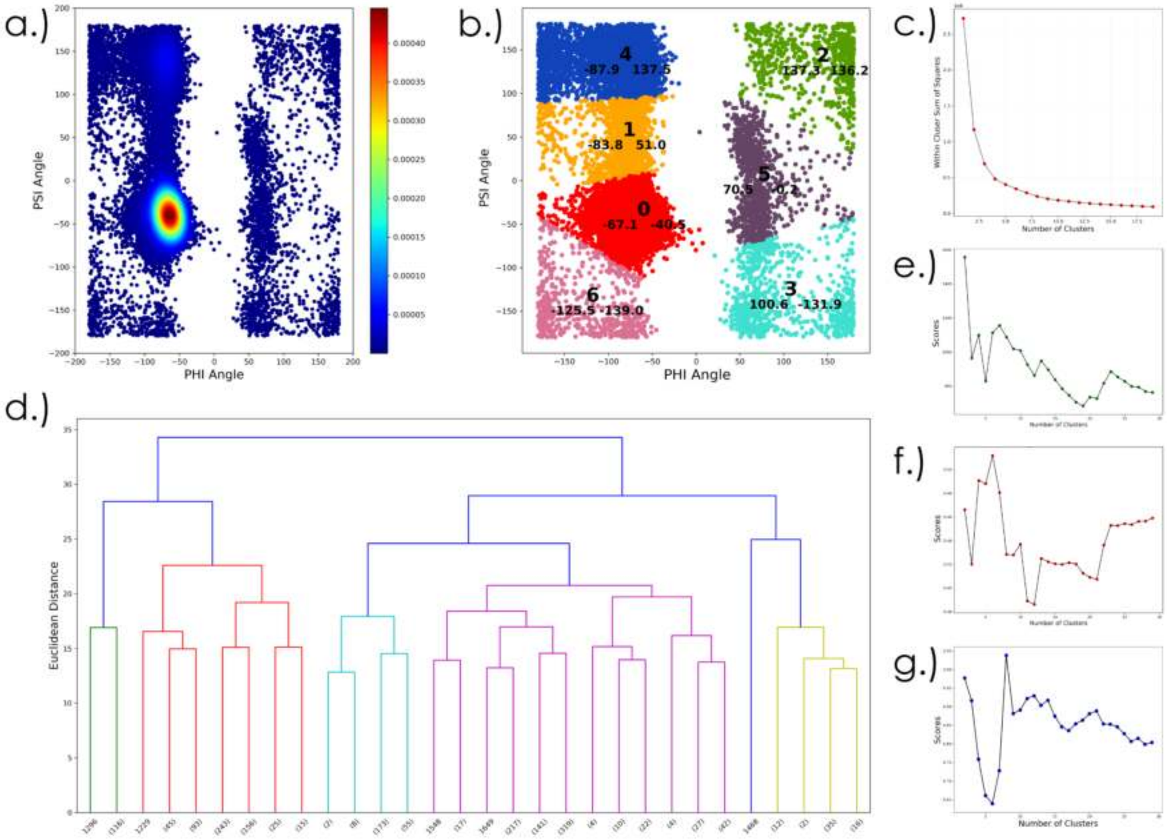


**Figure S10**. StELa clustering scheme. The Phi/Psi backbone angles are first plotted on the Ramachandran plot[8-10] (a) and clustered with a centroid based method. (b) The number of clusters was determined using the within cluster sum of squares method. (c) as well as by considering the separation of groups to distinguish between the alpha, beta and bridge regions.[11] The structure is described with the centroid labels and clustered with complete linkage hierarchical clustering. The number of clusters was determined using a variety of statistical methods including the dendrogram (d), the local maximum from the Calinski-Harabasz score method (e), the local maximum from the silhouette score (f), and the local minimum from the Davies-Bouldin index (g).

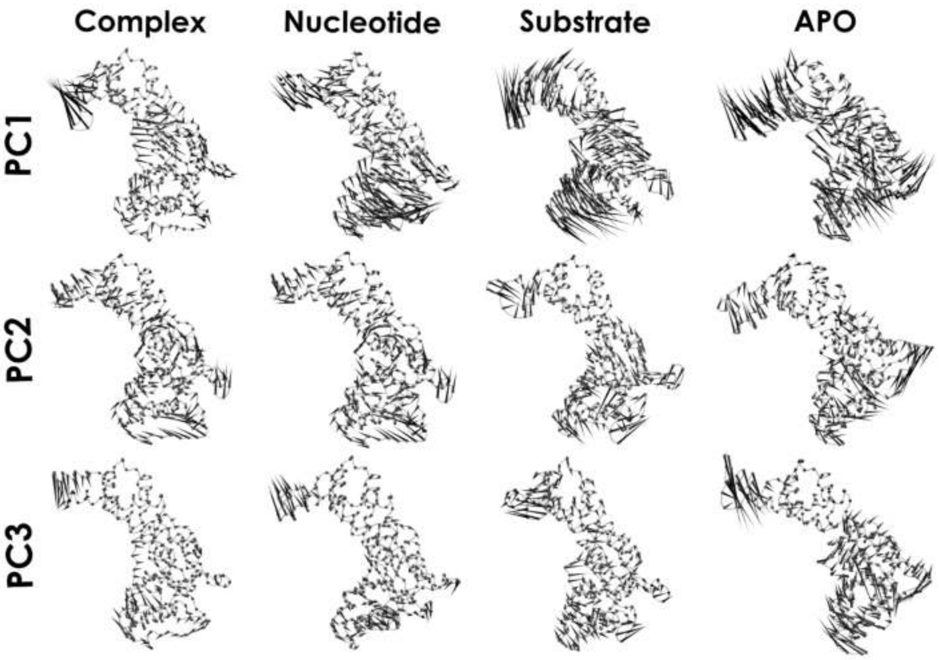


**Figure S11**. PC porcupine plots are similar to Figure 2, but for the katanin monomer setups, including the complex, nucleotide, substrate, and APO states. This and similar figures were produced using VMD.[12]

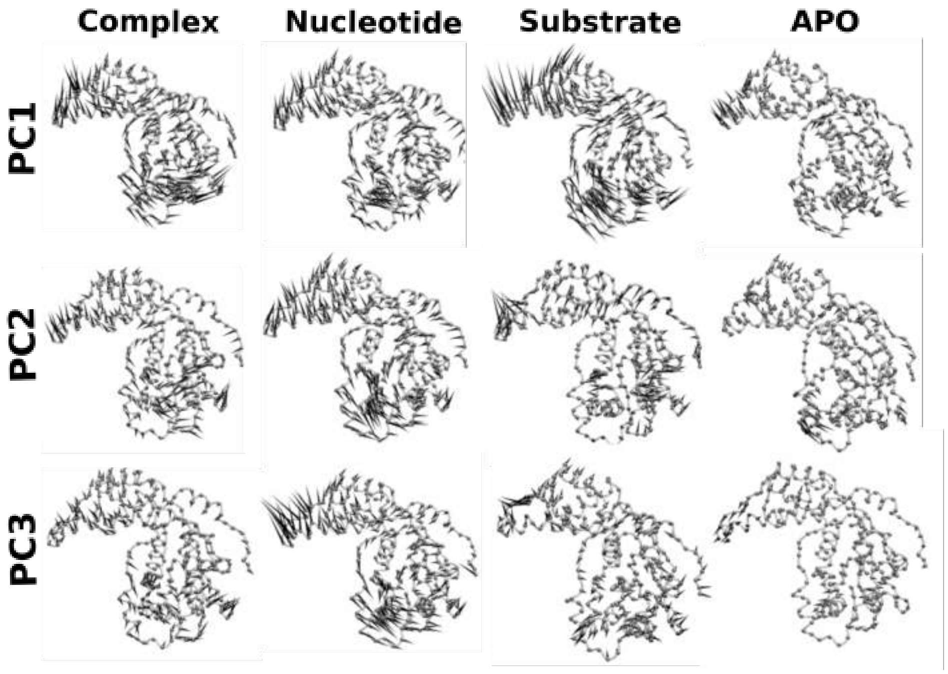


**Figure S12**. PC porcupine plots similar to Figure 2, but for the spastin monomer setups, including the complex, nucleotide, substrate, and APO states.

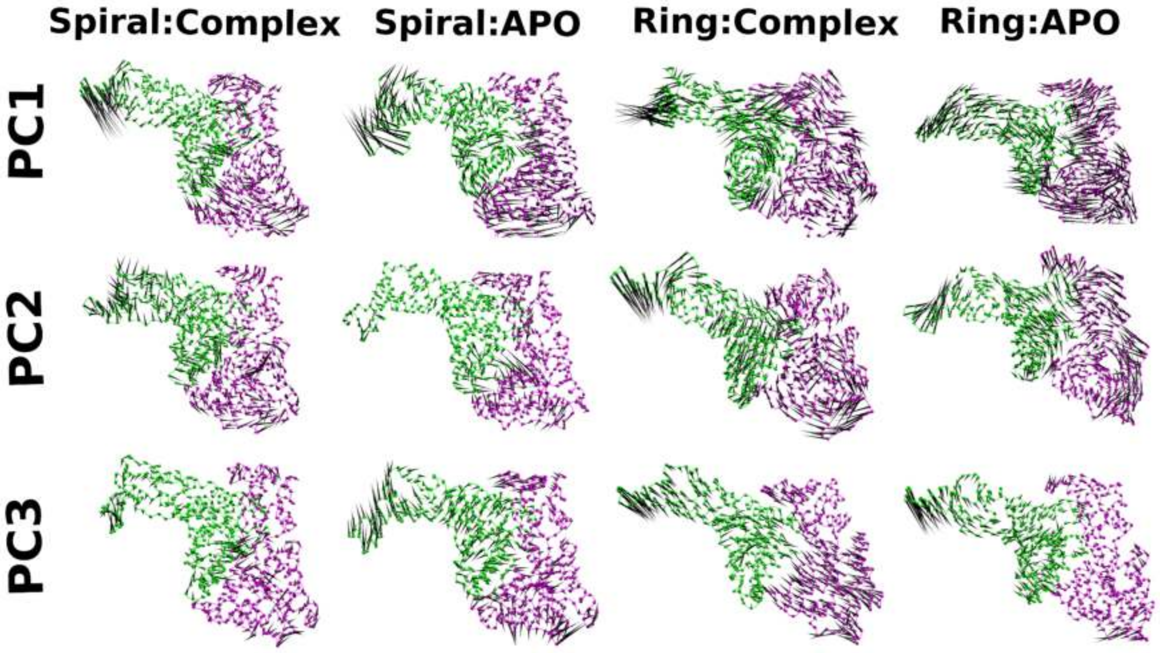


**Figure S13**: PC porcupine plots similar to Figure 2, but for the katanin AB-dimer setups.

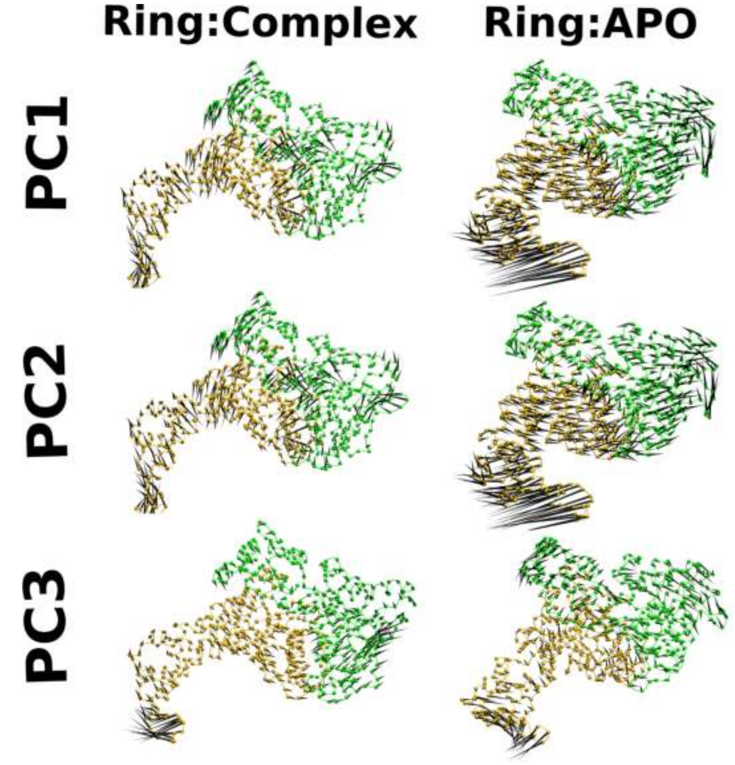


**Figure S14**: PC porcupine plots similar to Figure 2, but for the katanin BC-dimer setups.

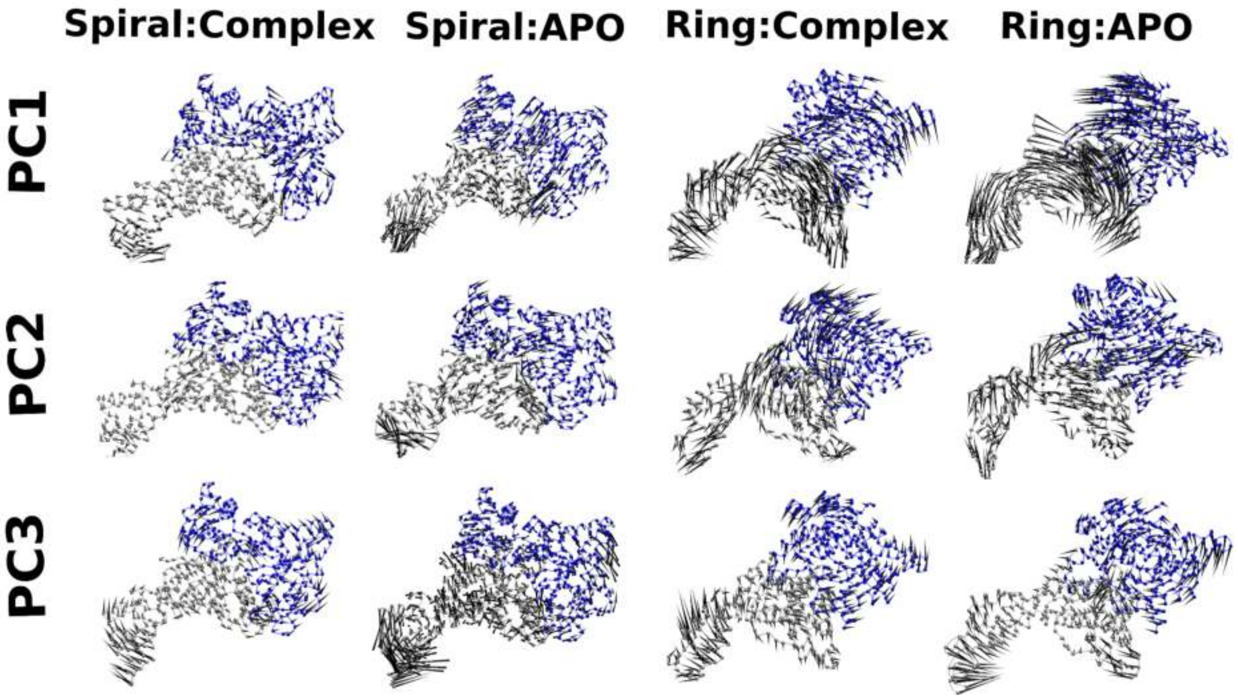


**Figure S15**: PC porcupine plots similar to Figure 3, but for the spastin EF-dimer setups.

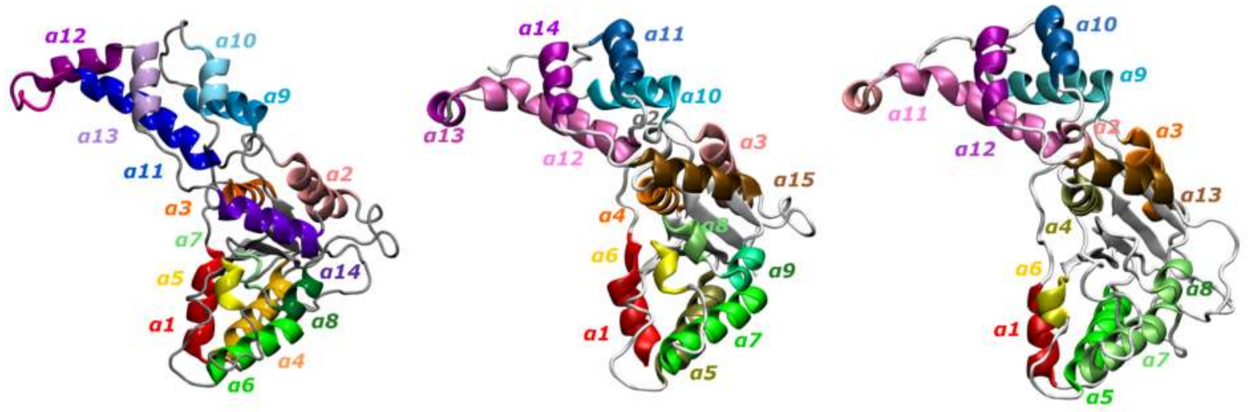


**Figure S16**: Hot Spots Analysis structure references to indicate which helices are which. (Left) The alpha helices are colored and labeled for the katanin spiral and ring (6UGD and 6UGE) monomer. (Middle) The alpha helices are colored and labeled for the spastin spiral (6P07) monomer. (Right) The alpha helices are colored and labeled for the spastin ring (6PEN) monomer.

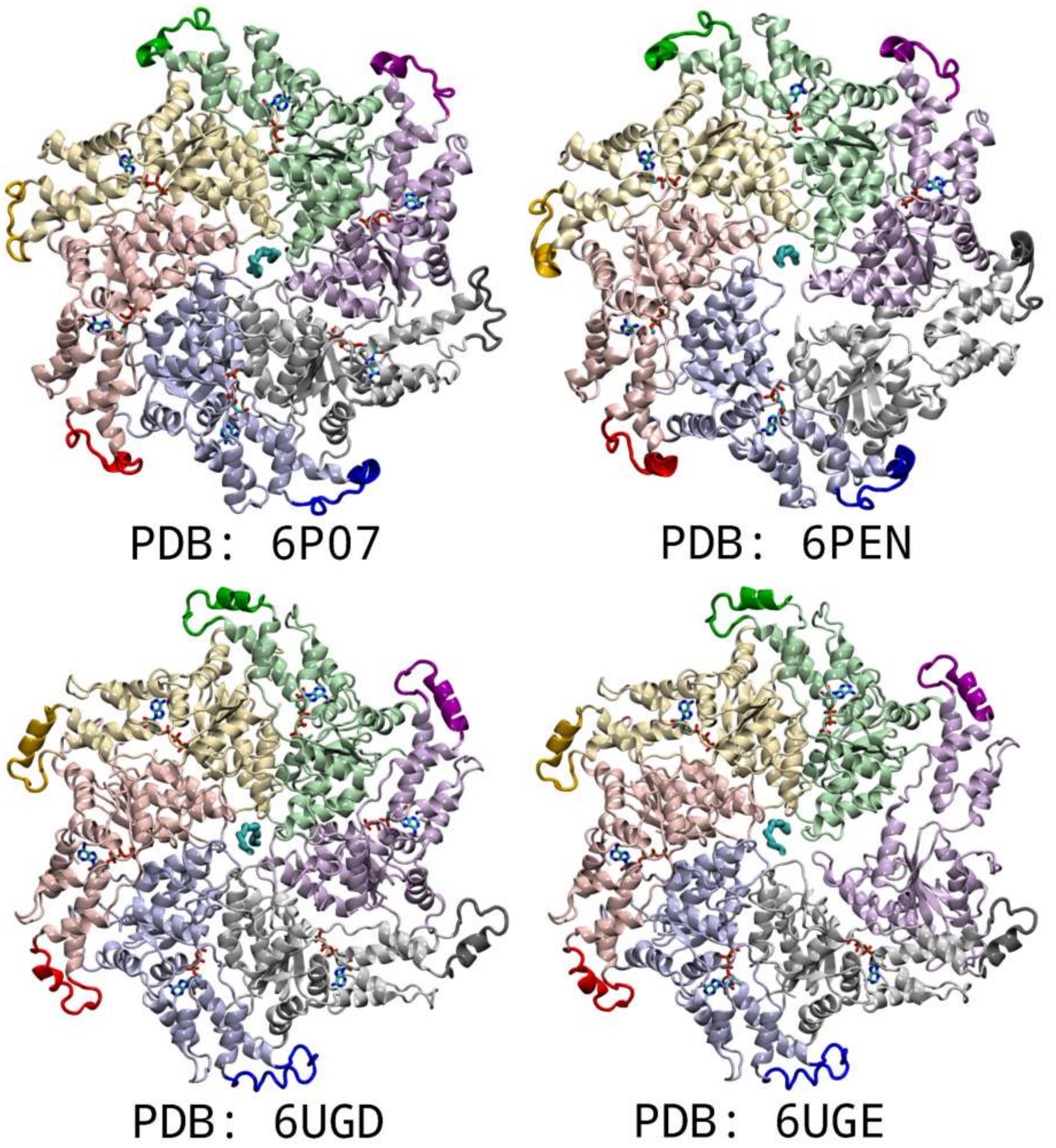


**Figure S17**: Starting configurations from the cryo-EM structures used in our MD simulations: spastin spiral (6P07), spastin ring (6PEN), katanin spiral (6UGD), and katanin ring (6UGE).[1-4] The hexamers are colored by monomer (A - purple - to F - grey). The monomers are shown in pastel colors while the HBD tip are bolded.

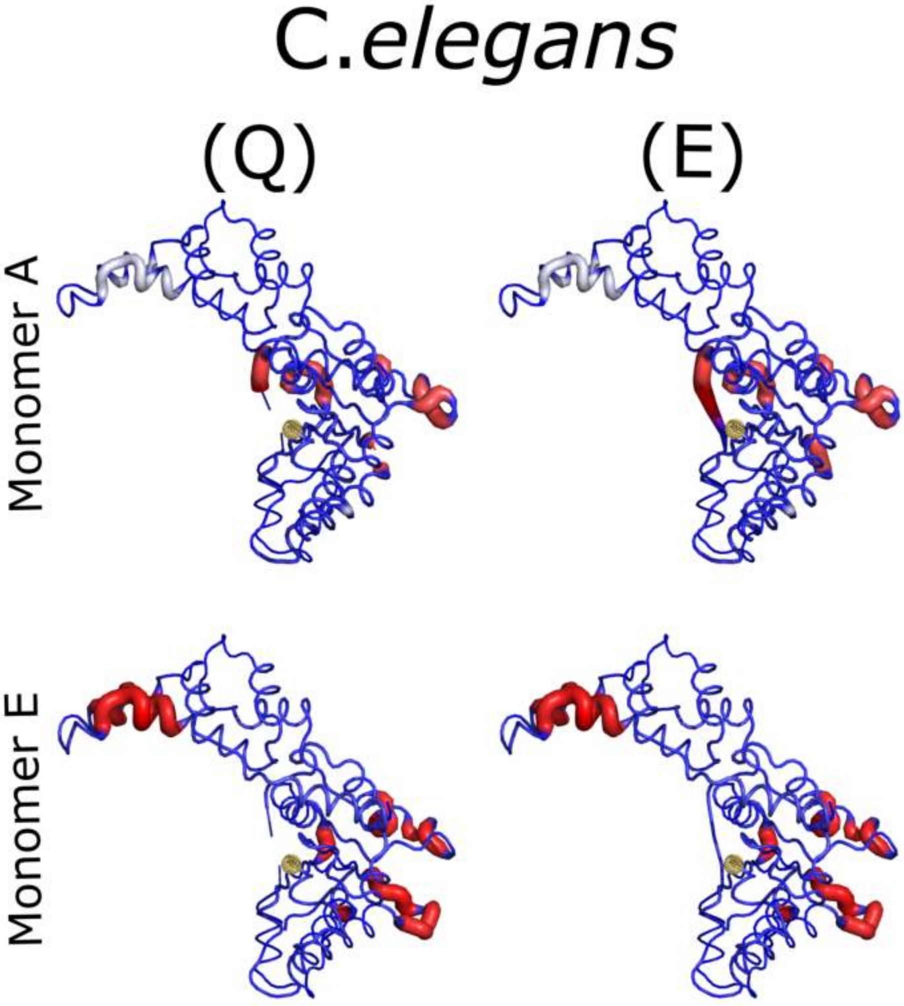


**Figure S1**8 DynaMut deformation energy profiles provide a measure over the first 10 non-trivial modes of the amount of local flexibility in the protein.[13] The regions highlighted in blue are considered stabilizing and the regions highlighted in red are considered destabilizing. Residue 293 for katanin from C.*elegans* (6UGD) was mutated from an E to a Q in the Cryo-EM and is highlighted as a gold sphere.[1,2] The structure of the HBD tip is noticeably different (as seen in **Figure S17**) in monomer A which is a Loop-Helix structure and monomer E which is completely unstructured. The deformation energies due to the mutation are not noticeably different, however, the energies due to the difference in the monomers were. The profiles indicate that this residue is allosterically connected to regions in the substrate binding region, the formation of the convex interface, and the HBD tip. In monomer E, these energies were found to be largely destabilized in comparison to monomer A.

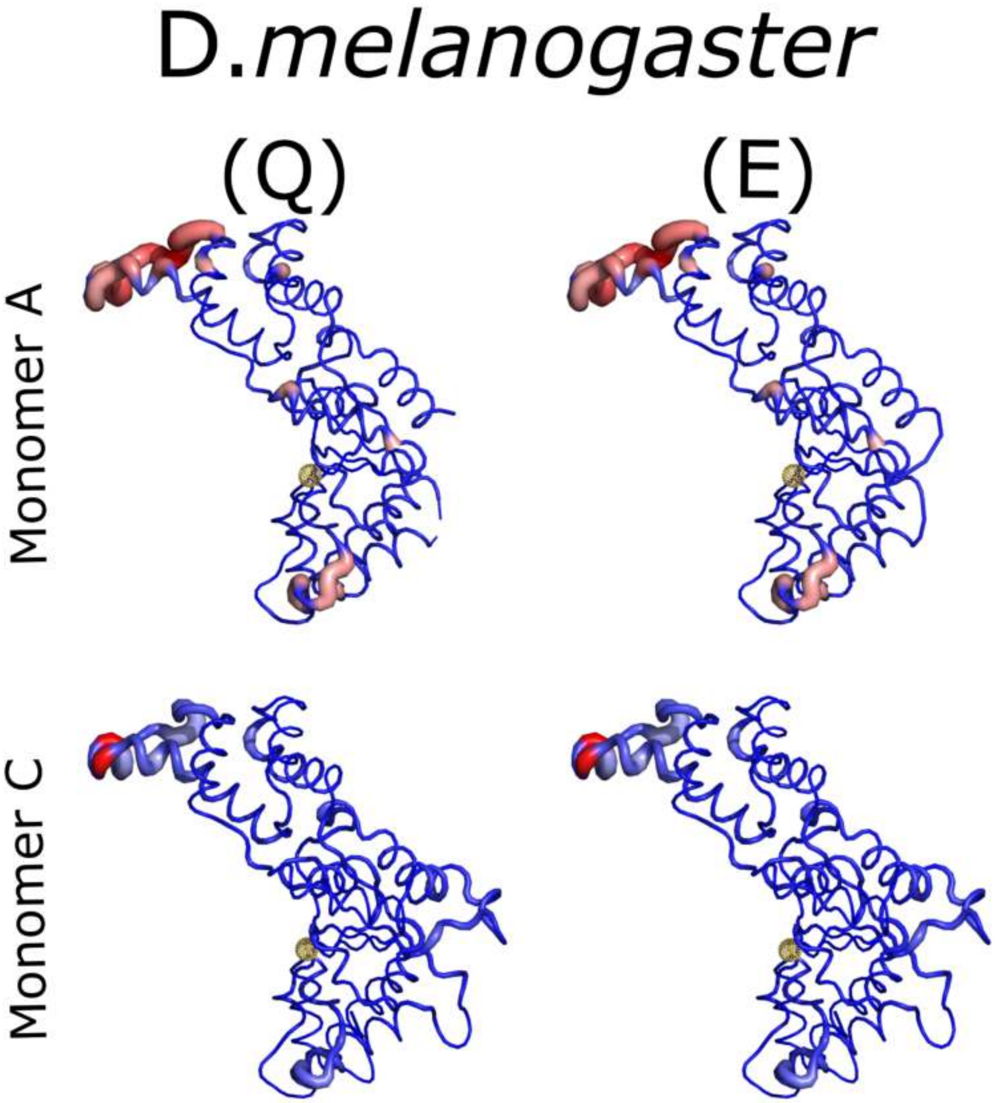


**Figure S19**: DynaMut deformation energy profiles provide a measure over the first 10 non-trivial modes for the amount of local flexibility in the protein.[13] The regions highlighted in blue are considered stabilizing and the regions highlighted in red are considered destabilizing. Residue 583 in spastin from D.*melanogaster* (6P07) was mutated from an E to a Q in the Cryo-EM structure and is highlighted as a gold sphere.[3] The structure of the HBD tip is noticeably different (as seen in **Figure S16**) in monomer A, which is a Helix-Loop structure, and monomer C, which is completely unstructured. The deformation energies due to the mutation are not noticeably different. However, the magnitude of the deformation energies differ between the two monomers. The HBD tip and substrate binding region in monomer A is largely destabilized in comparison to monomer C where these regions are stabilized.

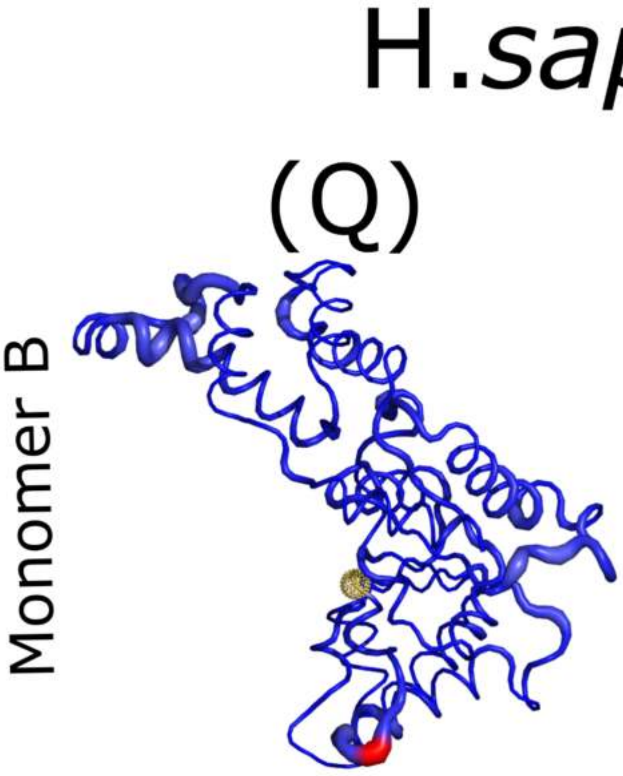


**Figure S20**: DynaMut deformation energy profiles provide a measure over the first 10 non-trivial modes for the amount of local flexibility in the protein.[13] The regions highlighted in blue are considered stabilizing and the regions highlighted in red are considered destabilizing. Residue 442 in spastin from H.*sapiens* (6PEN) was not mutated in the Cryo-EM and is highlighted as a gold sphere.[4] The structure of the HBD tip is the same in all monomers (as seen in **Figure S16**), which is a Helix-Loop structure. The deformation energies due to the mutation are not noticeably different. This residue is found to have an allosteric connection to the substrate binding region and the HBD tip.

| Species: Protein | Mutation | Chain | ΔΔG (kcal/mol) |
|---|---|---|---|
| C.*elegans* : Katanin | Q293E | A | -0.545 |
| | | E | 0.191 |
| D.*melanogaster* : Spastin | Q583E | A | -0.148 |
| | | C | -0.898 |
| H.*sapiens* : Spastin | E442Q | B | -0.434 |

**Table S7**. The outcome ΔΔG from the DynaMut single mutation predictions for the Walker B mutation.[13] In C.*elegans* katanin and D.*melanogaster* spastin, the Walker B was mutated from an E to a Q in the cryo-EM structure to prevent hydrolysis so in this analysis, the solved Q was mutated to the WT E.[1-3] In katanin, the mutation was dramatically destabilizing in monomer A, which exhibits the typical Loop-Helix structure for the HBD tip and interestingly slightly stabilizing in monomer E, which exhibits the Loop-Loop structure for the HBD tip. In spastin, both monomers experience a destabilizing effect although it is notably larger for monomer C, which adopts a Loop-Loop structure. In H.*sapiens* spastin, no Walker B mutation was present so the WT E was mutated to a Q which resulted in a destabilizing effect.[4] The positive values are considered stabilizing and the negative values are considered destabilizing.

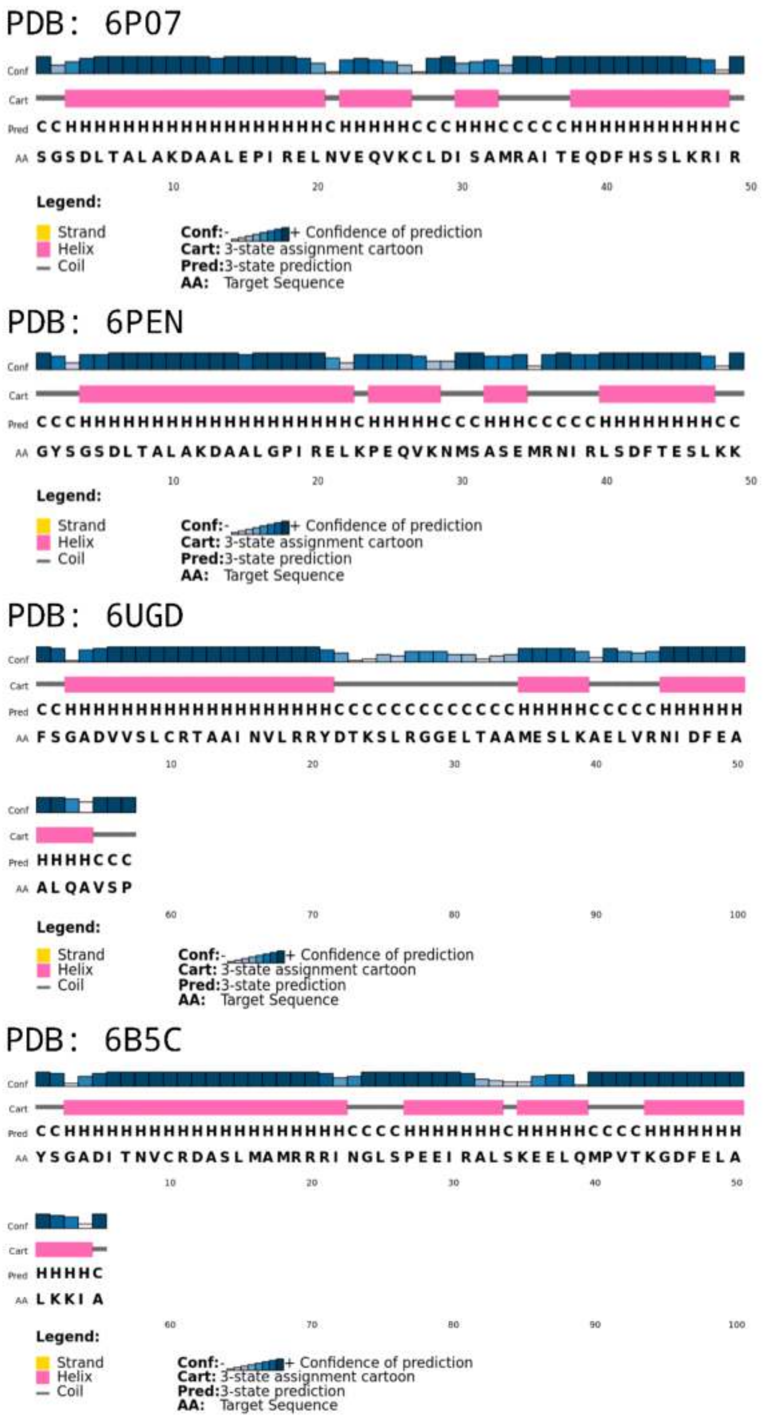


**Figure S21**: The PSIPRED results for prediction of secondary structure of the identified region, dubbed the HBD tip, based on the amino acid sequence for each indicated species of interest (Spastin - D.*melanogaster*-6P07, and H.*sapiens*-6PEN; Katanin - C.*elegans*-6UGD, and H.*sapiens*-6B5C).[14]

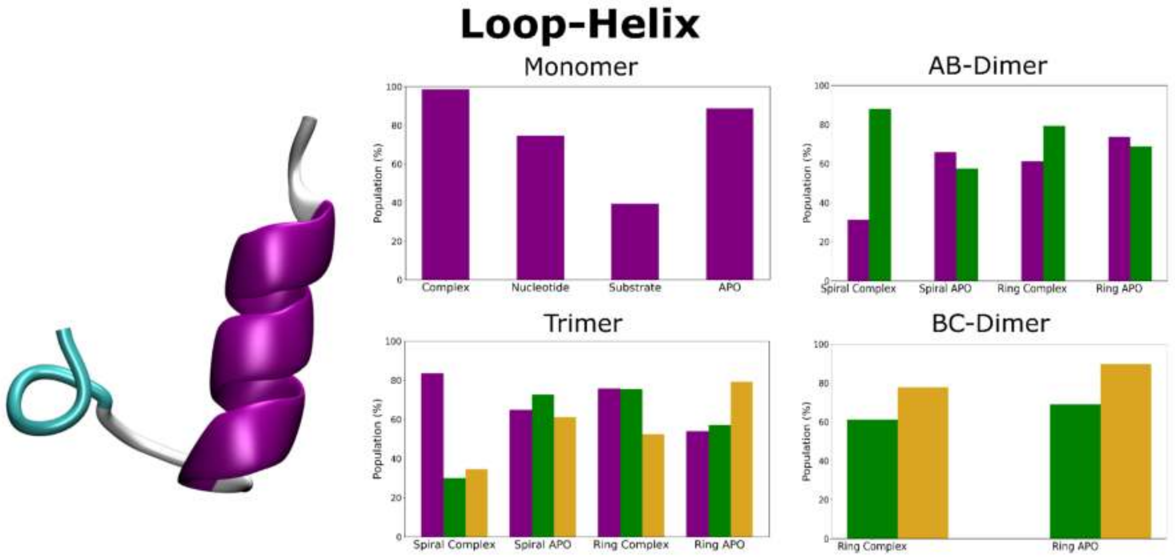


**Figure S22**: The population of states identified to have a Loop-Helix structure in all katanin systems, which was found to be the starting structure in simulation.[1,2] Monomer A is in purple, monomer B is in green, & monomer C is in yellow.

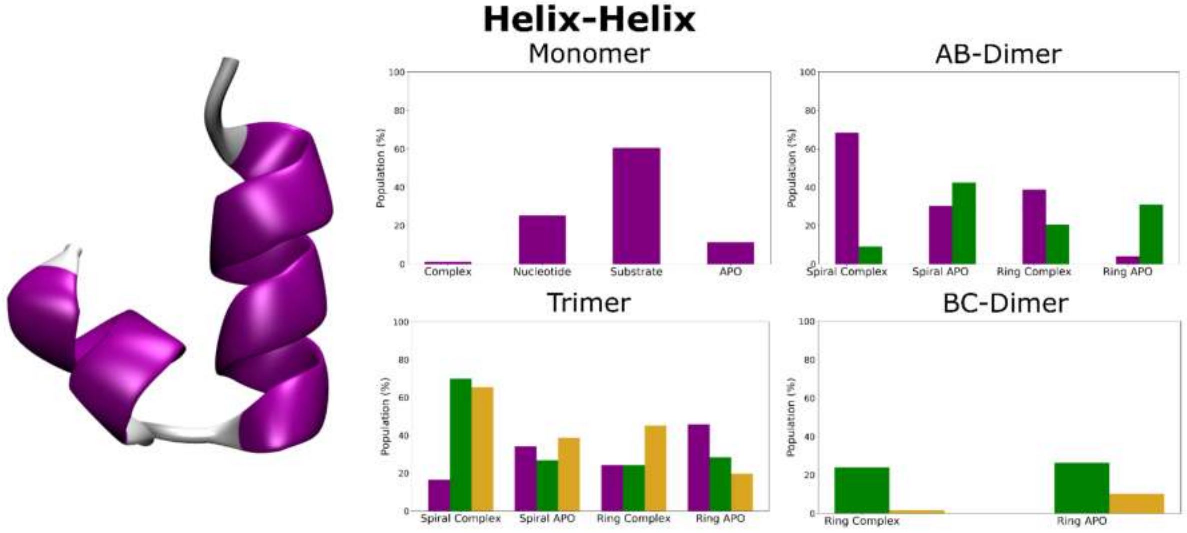


**Figure S23**: The population of states identified to have a Helix-Helix structure in all katanin systems. This structure was found to be in higher population in the presence of a single binding partner, but not both in the monomer. This structure is thought to be the preferred pathway to a 2 helix system and quite stable.[15] Monomer A is in purple, monomer B is in green, & monomer C is in yellow.

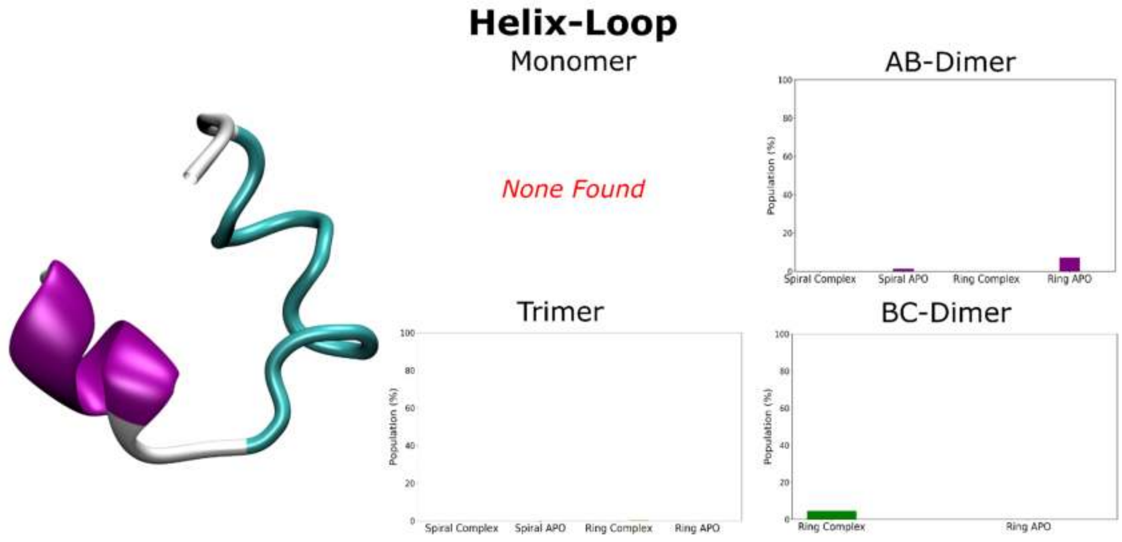


**Figure S24**: The population of structures identified to have a Helix-Loop structure in all katanin systems. This structure is one of the referenced flexible ensembles. Monomer A is in purple, monomer B is in green, & monomer C is in yellow. This structure was not present in the monomer and in the Trimer Ring Complex.

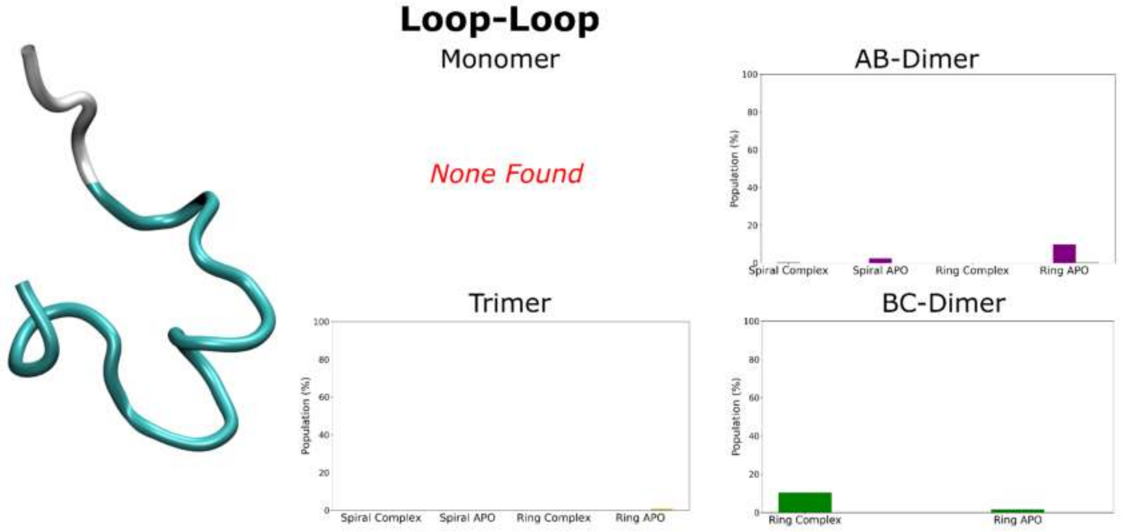


**Figure S25**: The population of structures identified to have a Loop-Loop structure in all katanin systems. This structure is one of the referenced flexible ensembles. Monomer A is in purple, monomer B is in green, and monomer C is in yellow. This structure was not present in the monomer, the AB Dimer Spiral APO, and the Trimer Ring APO.

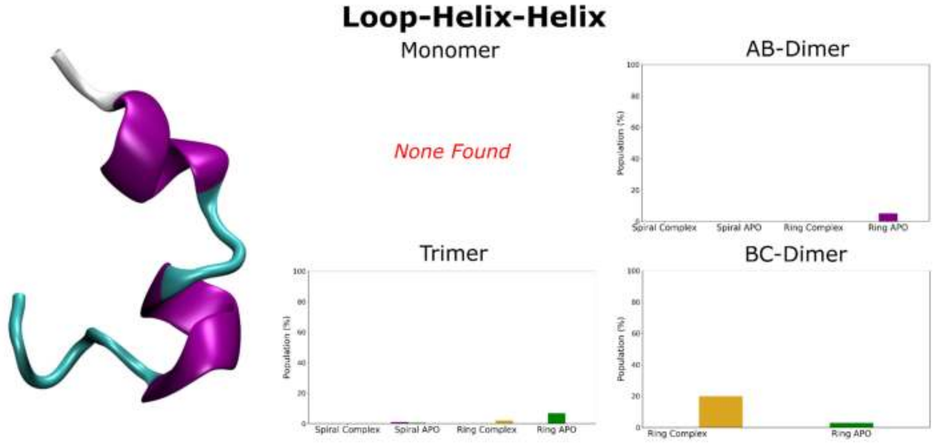


**Figure S26**: The population of structures identified to have a Loop-Helix-Helix structure in all katanin systems. This structure is of interest as it resembles the structure found in the H.*sapiens* solved structure which is described as the “nucleotide-free monomer”.[15] It is also striking that this structure is the result of the initial helix breaking in half to create two distinct helices. This structure is one of the referenced flexible ensembles. Monomer A is in purple, monomer B is in green, and monomer C is in yellow. This structure was not present in the monomer, the Trimer Spiral APO, and in the Ring Complex.

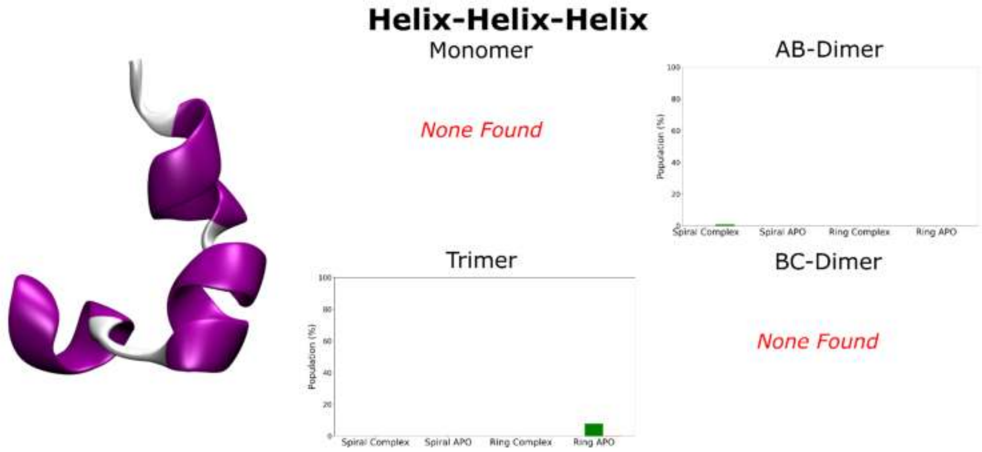


**Figure S27**: The population of structures identified to have a Helix-Helix-Helix structure in all katanin systems. This ensemble is particularly interesting as it experiences the loop to helix transition and the split helix structure. This structure is one of the referenced flexible ensembles. Monomer A is in purple, monomer B is in green, and monomer C is in yellow. This structure was not present in the monomer, the BC-Dimer, and in the Dimer Spiral Complex.

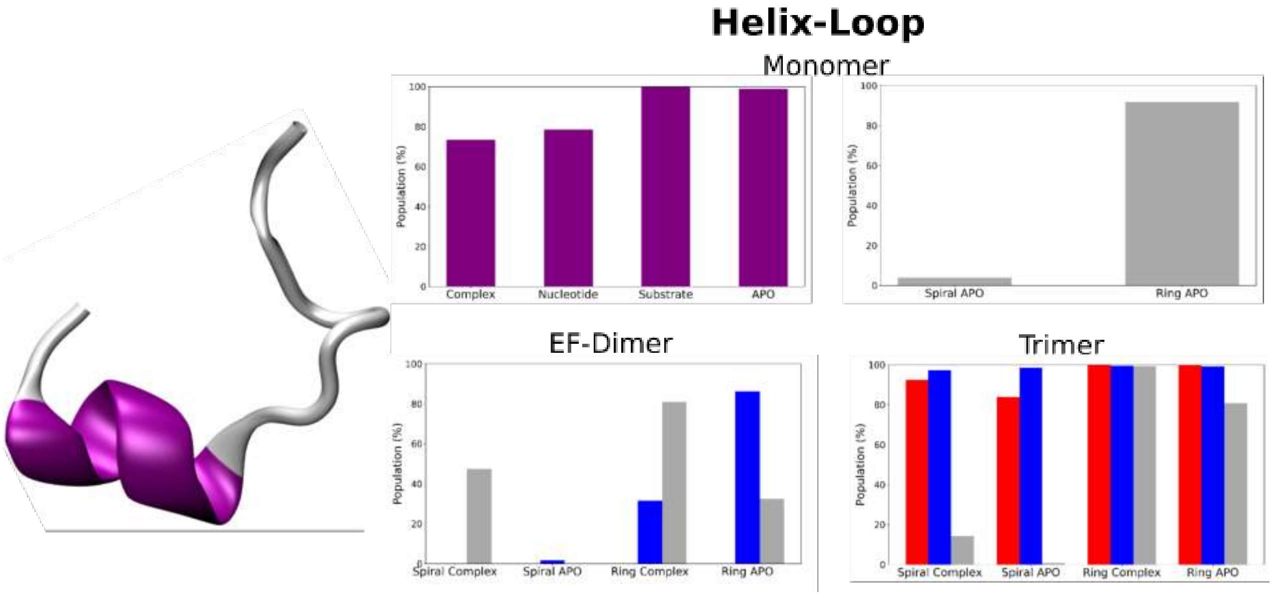


**Figure S28**: The population of structures identified to have a Helix-Loop structure in all spastin oligomers.[3,4] Monomer A is in purple, monomer D is in red, monomer E is in blue, and monomer F is in gray.

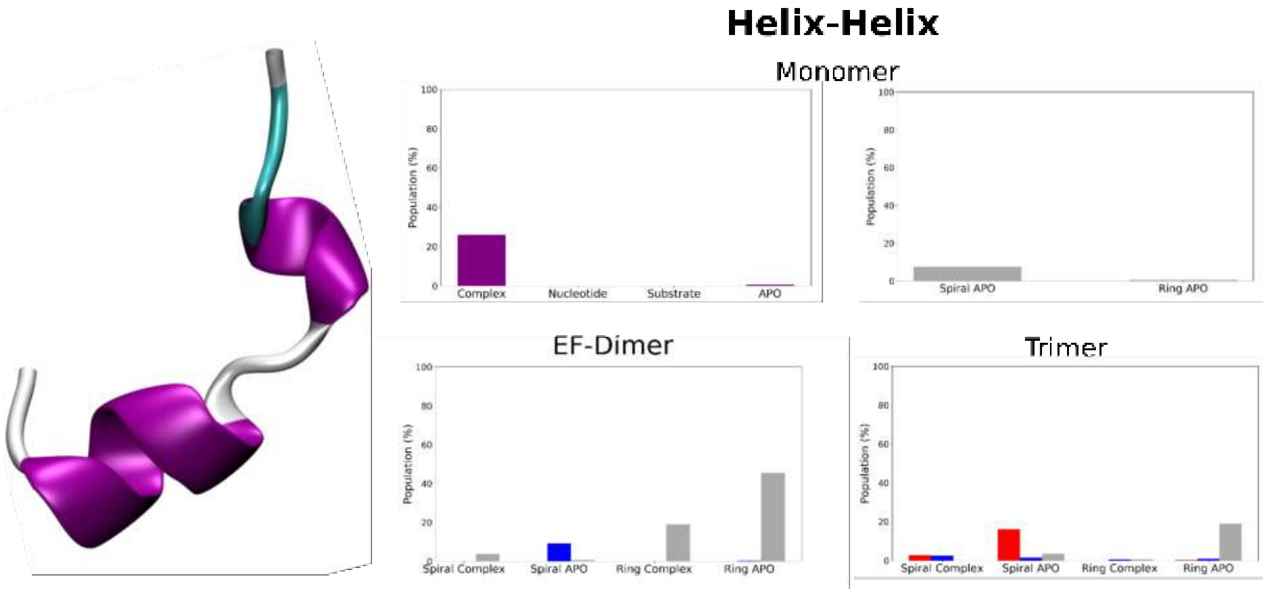


**Figure S29**: The population of structures identified to have a Helix-Helix structure in all spastin oligomers. Monomer A is in purple, monomer D is in red, monomer E is in blue, and monomer F is in gray.

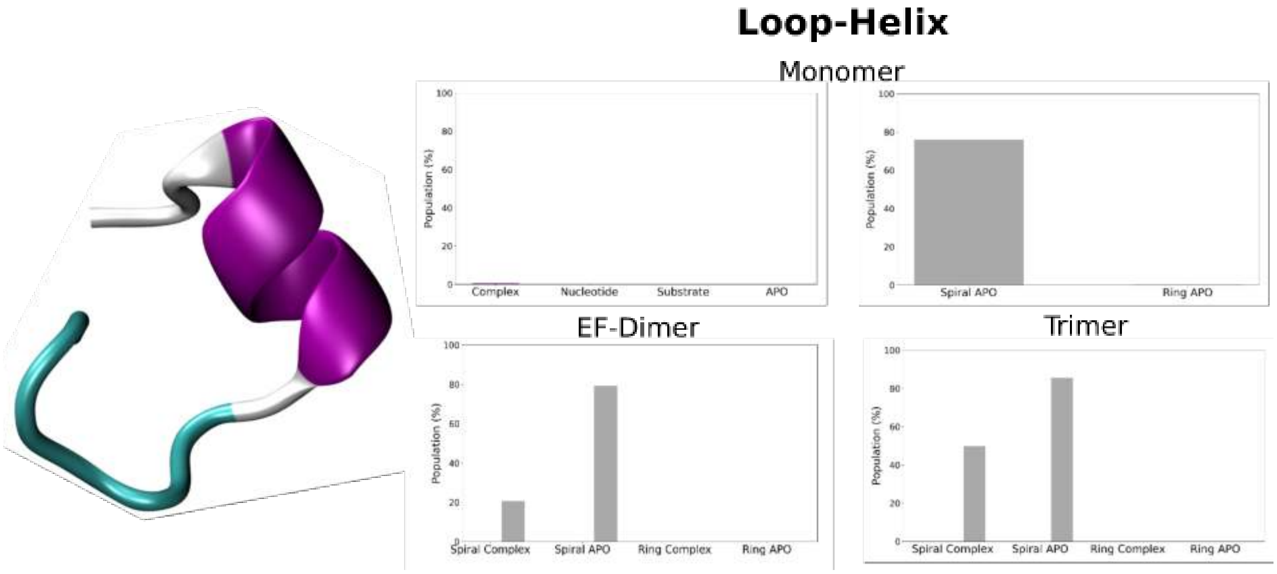


**Figure S30**: The population of structures identified to have a Loop-Helix structure in all spastin oligomers. Monomer A is in purple, monomer D is in red, monomer E is in blue, and monomer F is in gray. This structure was present in >1% in the complex monomer simulations.

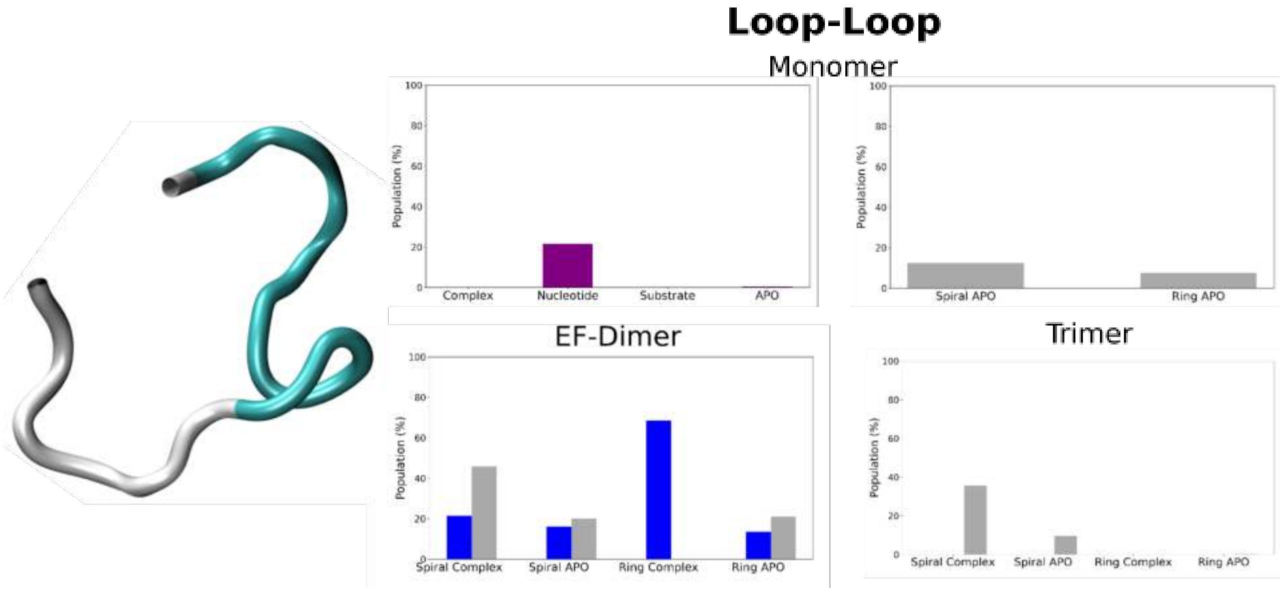


**Figure S31**: The population of structures identified to have a Loop-Loop structure in all spastin oligomers. Monomer A is in purple, monomer D is in red, monomer E is in blue, and monomer F is in gray.

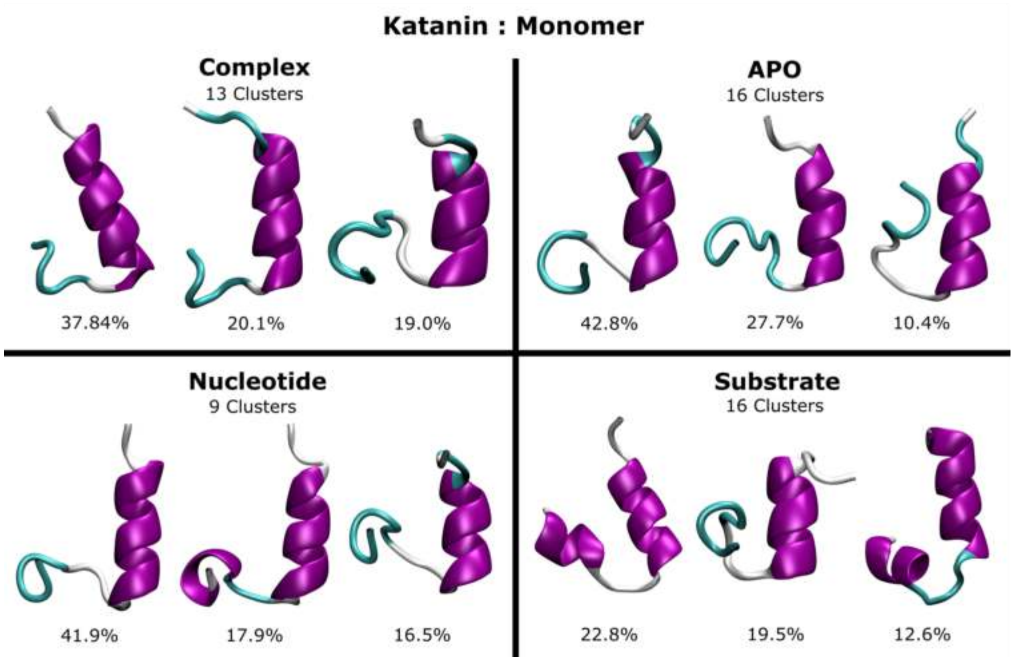


**Figure S32**: The representative structures of the top three populated clusters for the katanin monomer. The number of clusters used are indicated below the state and the population for the cluster is indicated below the respective cluster.

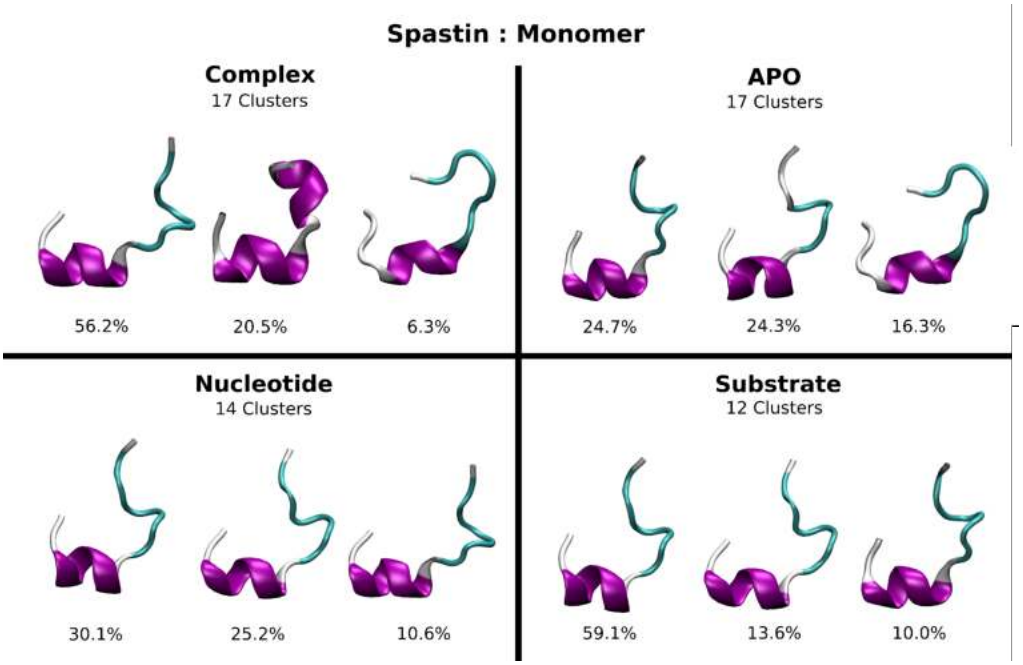


**Figure S33**: The representative structures of the top three populated clusters for the spastin monomer. The number of clusters used are indicated below the state and the population for the cluster is indicated below the respective cluster.
